# *In-Silico* Analyses of Molecular Force Sensors for Mechanical Characterization of Biological Systems

**DOI:** 10.1101/2024.07.17.603923

**Authors:** Diana M. Lopez, Carlos E. Castro, Marcos Sotomayor

## Abstract

Mechanical forces play key roles in biological processes such as cell migration and sensory perception. In recent years molecular force sensors have been developed as tools for *in situ* force measurements. Here we use all-atom steered molecular dynamics simulations to predict and study the relationship between design parameters and mechanical properties for three types of molecular force sensors commonly used in cellular biological research: two peptide-and one DNA-based. The peptide-based sensors consist of a pair of fluorescent proteins, which can undergo Förster resonance energy transfer (FRET), linked by spider silk (GPGGA)_n_ or synthetic (GGSGGS)_n_ disordered regions. The DNA-based sensor consists of two fluorophore-labeled strands of DNA that can be unzipped or sheared upon force application with a FRET signal as readout of dissociation. We simulated nine sensors, three of each kind. After equilibration, flexible peptide linkers of three different lengths were stretched by applying forces to their N-and C-terminal Cα atoms in opposite directions. Similarly, we equilibrated a DNA-based sensor and pulled on the phosphate atom of the terminal guanine of one strand and a selected phosphate atom on the other strand in the opposite direction. These simulations were performed at constant velocity (0.01 nm/ns – 10 nm/ns) and constant force (10 pN – 500 pN) for all versions of the sensors. Our results show how the force response of these sensors depends on their length, sequence, configuration and loading rate. Mechanistic insights gained from simulations analyses indicate that interpretation of experimental results should consider the influence of transient formation of secondary structure in peptide-based sensors and of overstretching in DNA-based sensors. These predictions can guide optimal fluorophore choice and facilitate the rational design of new sensors for use in protein, DNA, hybrid systems, and molecular devices.

**STATEMENT OF SIGNIFICANCE:** Biomolecular structures involved in various biological processes, including muscle function and sensory perception, generate, convey, and respond to mechanical forces. *In-vivo* accurate measurement of these forces is challenging but needed to understand biological function. Here we present a comprehensive computational analysis of three different types of molecular force sensors used to report pico-Newton level forces in biomolecular systems. Our atom-level simulation predictions provide mechanistic insight that can facilitate experimental data interpretation, selection of sensor design parameters, and the development of new force sensors tailored to specific applications and environments.

## INTRODUCTION

Biological processes such as receptor mediated signaling, cell migration, mechanotransduction, and muscle function often involve generation and detection of pico-to nano-Newton level forces (1–7). Single-molecule force spectroscopy techniques such as atomic force microscopy (AFM), optical tweezers, magnetic tweezers, and acoustic force spectroscopy are well suited to study force-induced structural changes in biomolecular structures *in vitro* (8–12). However, these techniques use perturbing external forces as well as micron-scale probes such as beads or functionalized cantilevers that can limit their ability to characterize biological processes *in situ*, for example in live cell assays.

Molecular force sensors have revolutionized the study of cellular mechanical responses, facilitating the *in-vivo* characterization of biomolecules like vinculin, integrin, and membrane receptors responding to intracellular and extracellular endogenous forces (13–21). Typically, these sensors are constructed by placing a flexible peptide tethered between a pair of fluorescent proteins that can undergo Förster resonance energy transfer (FRET) (Fig. 1 *A-C*) (22). The flexible protein fragment enables the response to force while the fluorescent proteins offer a mechanism for force readout. The distance between the fluorescent proteins is modified when applied forces stretch the peptide, thereby reducing FRET efficiency. Conversely, decreased force brings the fluorophores closer, thereby increasing FRET efficiency (13,23). In contrast, DNA-based sensors are constructed using a double stranded DNA duplex that ruptures at different forces, with the rupture event being the force readout. The DNA-based sensors function as binary reporters where the base-pairing between the two strands and the location of the force application points regulates the force threshold for rupture. Fluorescence signals from a single fluorophore, or a fluorophore with a quencher, can also be used as readouts to detect DNA dissociation under force. Any molecular event exerting sufficient force to rupture the duplex will trigger dissociation of the two strands leading to loss of fluorescence due to the fluorophore leaving the imaging plane (22,24) or increased fluorescence in the case of a fluorophore-quencher pair (18).

**Figure 1.**
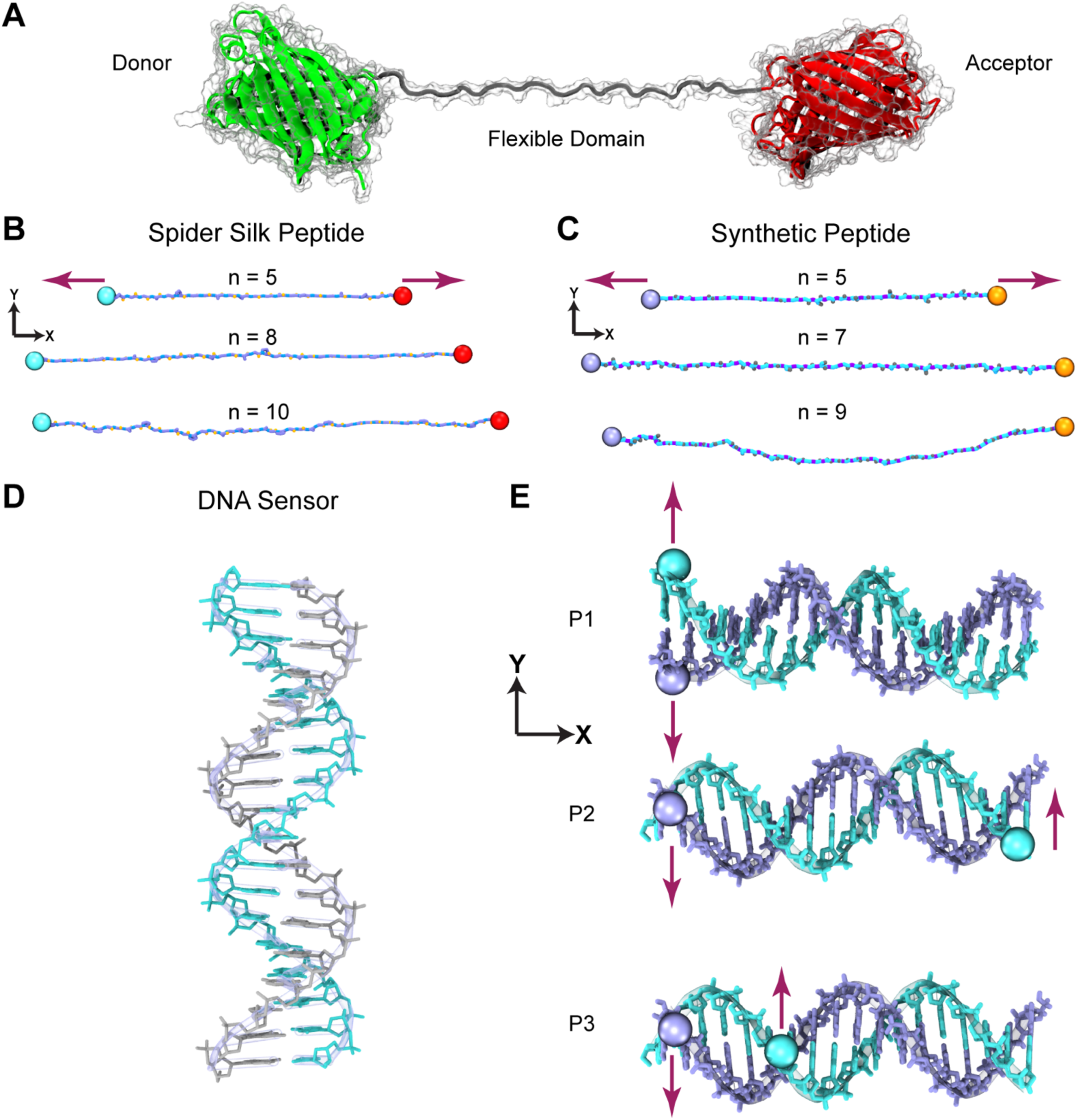
Molecular representations of peptide-and DNA-based sensors. (*A*) Molecular representation of a FRET-peptide force sensor. (*B*) Atomistic representation of a disorder region derived from the spider silk flagelliform protein (GPGGA)_n_ at different peptide lengths (n). (*C*) Atomistic representation of the synthetic versions of spider silk (GGSGGS)_n_ shown as in (*B*). Spheres denote stretched atoms during simulations and red arrows represent the direction of the forces applied in the SMD simulations. (*D*) Atomistic representation of an 18 bp DNA sensor. (*E*) Stretching positions P1 (unzipping), P2 (shearing), and P3 (mix) used in SMD simulations and experiments.

The most commonly used sequences for the elastic peptide domains of force sensors are derived from spider silk, which has been of interest for industrial and biological applications due to its high strength and ability to rapidly stretch and contract (25–27). Individual peptide units of silk proteins like flagelliform have been used to develop biomolecular sensors including FRET-based force sensors (13,23). To expand and improve the applications of spider silk peptides and synthetic derivatives it is critical to understand and probe their mechanical properties and how these depend on molecular parameters and loading rates. To date, a few studies have tested the effects of design parameters like length and composition of peptide-based sensor linkers computationally or experimentally (28–31). Some of these studies have revealed the mechanical effects of flow dependent stretching of spider silk, the influence of polyglycine regions and external forces on the elasticity of spider silk peptides, and details of fiber assembly. However, further studies that integrate the relationship between atom-level structure and dynamics, sequence, elasticity, and readout choice as design parameters could guide the design of future force sensors for tailored sensing responses.

Similarly, a few studies (32–34) have explored the mechanics of DNA-based sensors. Theoretical modeling and coarse-grained simulations of DNA duplexes indicate that rupture force thresholds depend on loading rate, time scale of observation, and pulling configuration. Notably, these models have not explored some aspects of the force-extension behavior, for example the formation of stretched states of DNA (S-DNA). An all-atom simulation study also pointed out the influence of bound drug molecules on the stability of short DNA duplexes stretched at fast velocities (33). Understanding DNA-based force sensor dynamics and mechanical responses at an atomistic level can expand the options for force sensor design as these provide a binary response and the option of multiple force readouts utilizing only a single DNA sequence of fixed length (32,33).

Here we present results from all-atom molecular dynamics (MD) simulations (35) characterizing the mechanical properties of three types of molecular force sensors typically used in cellular and biochemical research (23,36,37): spider silk-based protein sensors—which consists of a fluorescent protein FRET pair linked by a spider silk disordered region derived from the flagelliform protein (GPGGA)_n_; synthetic-based protein sensors—also consisting of a fluorescent protein FRET pair linked by a disordered synthetic region (GGSGGS)_n_ (23), and DNA-based sensors—consisting of a double stranded DNA region that can be unzipped or sheared upon stretching, depending on the designed location of force application (22,34).

Structural models of peptide linkers and DNA sensors were solvated, energy minimized, and equilibrated under zero force. Steered MD (SMD) (38) at constant velocity and constant force were used to predict the mechanical characteristics of these sensors. We found that predicted spring constants for peptide-based sensors stretched at constant velocities in the range of 0.01 nm/ns – 1 nm/ns or at constant forces in the range of 10 pN-50 pN are in general agreement with experiments (36). We also report potentially important effects of transient secondary structure formation, which could be important considerations for sensor designs, and the impact that selecting protein FRET pairs with different Förster radii (39) may have on observed sensor characteristics. Our simulations predict that shearing of a DNA-based sensor requires larger forces for shearing than unzipping as expected from experiments and give useful insight into the sensor design features that govern the rupture forces (22). We also study a region of DNA duplex stretching that might be used for a non-binary DNA-based sensor design. Overall, these results offer valuable insights into running simulations for effective comparisons with experiments and establish a framework for future computational studies and rational design of molecular force sensors.

## METHODS

### Modeling

To build the all-atom models of each of the six peptide variations for (GPGGA)_n_ (n = 5, 8, 10) and (GGSGGS)_n_ (n = 5, 7 and 9; Fig. 1), each amino acid was manually added and linked according to each peptide sequence using the modeling software COOT (40). For the dsDNA-based sensor we used the sequence from a previously reported DNA sensor (GGC CCG CAG CGA CCA CCC) (22) and used the web server “model.it” to generate the initial models (41). Hydrogen atoms were added using the psfgen plugin from visual molecular dynamics (VMD) (42). Water box generation and solvation of all nine systems was done with the solvate plugin from VMD. All systems were ionized with 200 mM NaCl and 1 mM MgCl_2_ using the autoionize plugin from VMD.

### MD Simulations

All MD simulations in this study were performed using NAMD 2.13 (43) with CHARMM36 (44) force field parameters for protein (peptide-based sensors) and nucleic acids (dsDNA-based sensors) in explicit TIP3P water. Simulations were conducted with a 2-fs integration step and a 12 Å cutoff for van der Waals interactions in the *NpT* ensemble at 300 K and 1 atm. All models underwent a 1,000-step energy minimization. This was followed by a 1 ns equilibration with a Langevin damping coefficient of γ = 1 ps^-1^, and then a 15 ns equilibration using γ = 0.1 ps^-1^. Spider silk and synthetic systems were initially stretched fully and naturally contracted during the minimization and equilibration stages.

Post-equilibration states served as the initial configurations for SMD simulations. The NAMD TCL forces interface was employed to do SMD at constant velocity. A virtual spring with a stiffness of *k* = 1 kcal mol^-1^ Å^-2^ was linked to the Cα of each terminus in spider silk and synthetic systems, and to a phosphorous atom on each strand of the dsDNA systems at designated anchor sites. For the peptide systems, these springs’ free ends were moved at constant velocity along the *x*-axis (corresponding to stretching the sensor), while the *y*-axis was chosen for the DNA systems (corresponding to dissociating the two strands). Simulation velocities were set to 0.1 nm/ns, 1 nm/ns, and 10 nm/ns for all systems, with and additional set of simulations at 0.01 nm/ns for peptide-based sensors. SMD simulations were also performed using constant forces ranging from 10 pN to 500 pN applied at the aforementioned anchor points and along the specified axes.

### Data Analysis

Theoretical spring constants (*k*_s_) for each peptide-based sensor type (spider silk and synthetic; Table 1) were computed using:

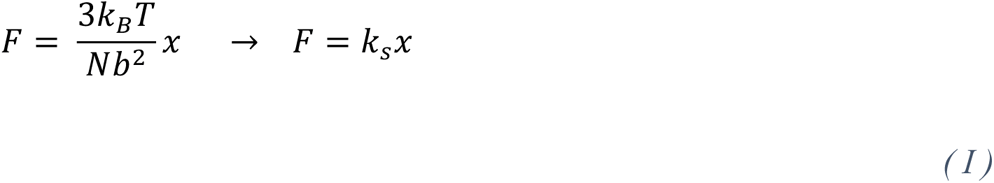

**Table 1.**
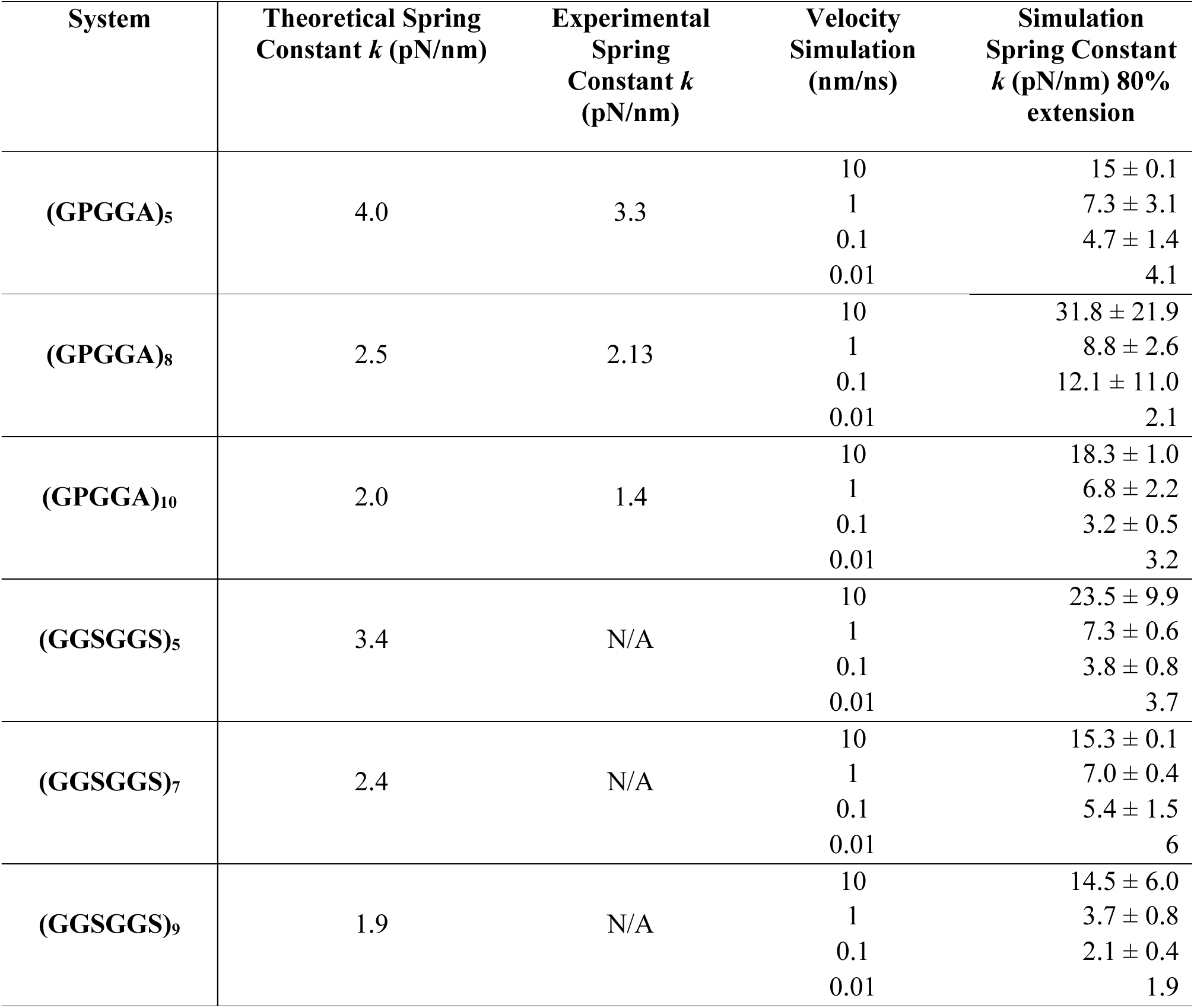
Spring constants of (GPGGA)_n_ and (GGSGGS)_n_ sensors from theory, experiments. (**36**)**, and simulations**

| System | Theoretical Spring Constant $k$ (pN/nm) | Experimental Spring Constant $k$ (pN/nm) | Velocity Simulation (nm/ns) | Simulation Spring Constant $k$ (pN/nm) 80% extension |
| --- | --- | --- | --- | --- |
| (GPGGA)<sub>5</sub> | 4.0 | 3.3 | 10 | 15 ± 0.1 |
|  |  |  | 1 | 7.3 ± 3.1 |
|  |  |  | 0.1 | 4.7 ± 1.4 |
|  |  |  | 0.01 | 4.1 |
| (GPGGA)<sub>8</sub> | 2.5 | 2.13 | 10 | 31.8 ± 21.9 |
|  |  |  | 1 | 8.8 ± 2.6 |
|  |  |  | 0.1 | 12.1 ± 11.0 |
|  |  |  | 0.01 | 2.1 |
| (GPGGA)<sub>10</sub> | 2.0 | 1.4 | 10 | 18.3 ± 1.0 |
|  |  |  | 1 | 6.8 ± 2.2 |
|  |  |  | 0.1 | 3.2 ± 0.5 |
|  |  |  | 0.01 | 3.2 |
| (GGSGGS)<sub>5</sub> | 3.4 | N/A | 10 | 23.5 ± 9.9 |
|  |  |  | 1 | 7.3 ± 0.6 |
|  |  |  | 0.1 | 3.8 ± 0.8 |
|  |  |  | 0.01 | 3.7 |
| (GGSGGS)<sub>7</sub> | 2.4 | N/A | 10 | 15.3 ± 0.1 |
|  |  |  | 1 | 7.0 ± 0.4 |
|  |  |  | 0.1 | 5.4 ± 1.5 |
|  |  |  | 0.01 | 6 |
| (GGSGGS)<sub>9</sub> | 1.9 | N/A | 10 | 14.5 ± 6.0 |
|  |  |  | 1 | 3.7 ± 0.8 |
|  |  |  | 0.1 | 2.1 ± 0.4 |
|  |  |  | 0.01 | 1.9 |

This equation, derived from the freely jointed chain (FJC) model, describes the polymer extension *x* of an entropic spring in the presence of a force *F*. The number of segments is *N*, the segment or Kuhn length is *b* (0.35 nm per amino acid), and *k_B_T* is the Boltzmann’s constant times absolute temperature (∼ 4.1 pN·nm at room temperature, 298 K).

Experimental spring constants of spider silk peptides (Table 1) were extracted from reported force extension curves obtained using optical tweezers (36).

All simulation trajectories were viewed in VMD to confirm full extension of peptides or rupture of DNA strands and to ensure that interactions across the periodic box did not occur. Force-extension data were extracted from output files using custom scripts and processed with the Xmgrace software. In constant-velocity SMD simulations of peptide-based sensors, force-extension curves were examined over extensions accounting for 80% of the theoretical extended length (N·0.35 nm) to exclude backbone overstretch and primarily describe the stiffness due to entropic elasticity. We also studied the stretched region of force-extension curves pertaining to FRET-relevant distances. A linear fit was performed to compute the slope representing the effective spring constant for each system while enforcing a value of zero force at the beginning of the stretching simulation (zero extension).

Trajectories resulting from constant-force SMD simulations were analyzed by fitting extension vs. time curves using:

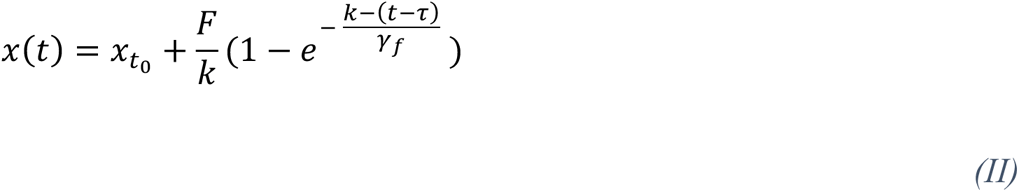

where *x* is the extension of an overdamped spring (extension of the linker), *t* represents time, *x*_*t0*_ is the extension of the linker at *t* = 0, *F* is force, *k* is the spring constant, and γ_f_ is the drag coefficient.

Secondary structure analysis of the SMD simulations of all peptide-based linkers was carried out using the timeline VMD plugin. Percentage of secondary structure present in each peptide for each trajectory was computed using customized scripts.

The radius of gyration was calculated using:

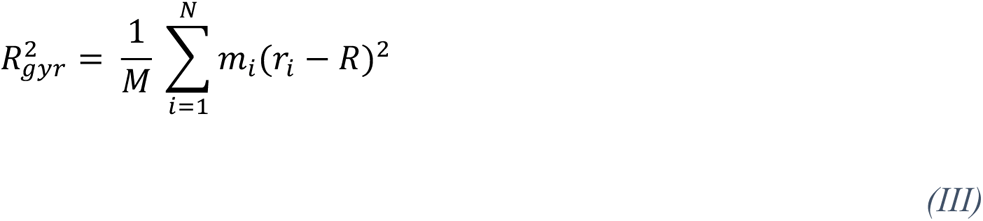

where *M* is the total mass of the peptide, *m_i_* is atomic mass, *R* is the center of mass of the peptide, *r_i_* is the center of mass of the atom, and *N* is the number of atoms. All peptides’ radii of gyration were computed throughout all trajectories using TCL scripts. VMD was used to render all the images and Videos of simulations.

## RESULTS

Force sensors are typically incorporated in series with biomolecular structures subjected to mechanical stimuli. In this way, sensors can report on the magnitude of the endogenous forces experienced by these systems *in situ.* Endogenous mechanical stimuli can be diverse – in some cases biomolecular structures will be subjected to rapid and sustained increases in force, while in others a constant force might be expected, with more complex time-dependent force stimuli depending on biological context. To explore the mechanical properties of peptide-and DNA-based sensors we carried out equilibrations followed by SMD simulations at both constant velocity and constant force.

### Elasticity of spider silk and synthetic linkers stretched at constant velocity

All-atom models of peptide-based force sensors in water and ions, including three spider silk ((GPGGA)_5_; (GPGGA)_8_; GPGGA)_10_) and three synthetic linkers ((GGSGGS)_5_; (GGSGGS)_7_; (GGSGGS)_9_), were equilibrated for 15 ns (Tables S1-S2; Video S1). The extended peptides quickly adopted a compact, disordered conformation. This was reflected by a decrease in the radius of gyration *R_gyr_* for most systems occurring over 2 ns to 8 ns, with *R_gyr_* values stabilizing shortly thereafter (Fig. S1). Although compact conformations for peptides were dynamic and disordered, we assumed that lack of changes in *R_gyr_* reflected equilibrated systems and used conformations at 10 ns and 15 ns as two different starting points for constant-velocity SMD simulations. These simulations were carried out at pulling velocities of 0.1 nm/ns, 1 nm/ns, and 10 nm/ns in duplicates (starting from the two different conformations obtained after 10 ns and 15 ns of equilibration) with an additional single 0.01 nm/ns SMD simulation for each of the six peptide-based sensor systems (Videos S2-3). In all cases we observed extension of the peptide, consistent with an increase in *R_gyr_* (Fig. S2).

Force-extension curves for the smallest spider silk system (GPGGA)_5_ were similar at all stretching velocities (Fig. 2 *A* and S3 *A*), with a first phase in which force increased linearly for extension values between ∼ 2 nm and ∼ 7 nm, followed by a second phase of rapid force increase for extension values between ∼ 7 nm and ∼ 9.5 nm. The first phase corresponded to peptide straightening where the response is largely mediated by entropic elasticity, while the second phase involved stretching of the peptide backbone, which occurs at forces that are beyond the typical range of physiological conditions. We performed linear regressions assuming zero force at zero extension, as expected from the FJC model (equation 1) and the setup of the SMD simulations. The resulting slopes of the force-extension curves during the first straightening phase of extension were used to estimate spring constants at each velocity (Fig. S3 and S4). We used data up to extensions that were 80% of the fully extended but not stretched peptide. These fits yielded (GPGGA)_5_ spring constants of (*k*_0.01_ = 4.1 pN/nm, *k*_0.1_ = 5.7 pN/nm, *k*_1_ = 9.4 pN/m, and *k*_10_ = 15.1 pN/nm) (results from individual simulations in Table S1) These results, especially for the 0.01 and 0.1 nm/ns speeds, agree well with the theoretical entropic spring constant of a FJC of the same length (4.03 pN/nm) and to the experimentally measured value for this peptide linker (3.3 pN/nm; Table 1) (36). We observed that the initial configuration had a small impact on the predicted stiffnesses.

**Figure 2.**
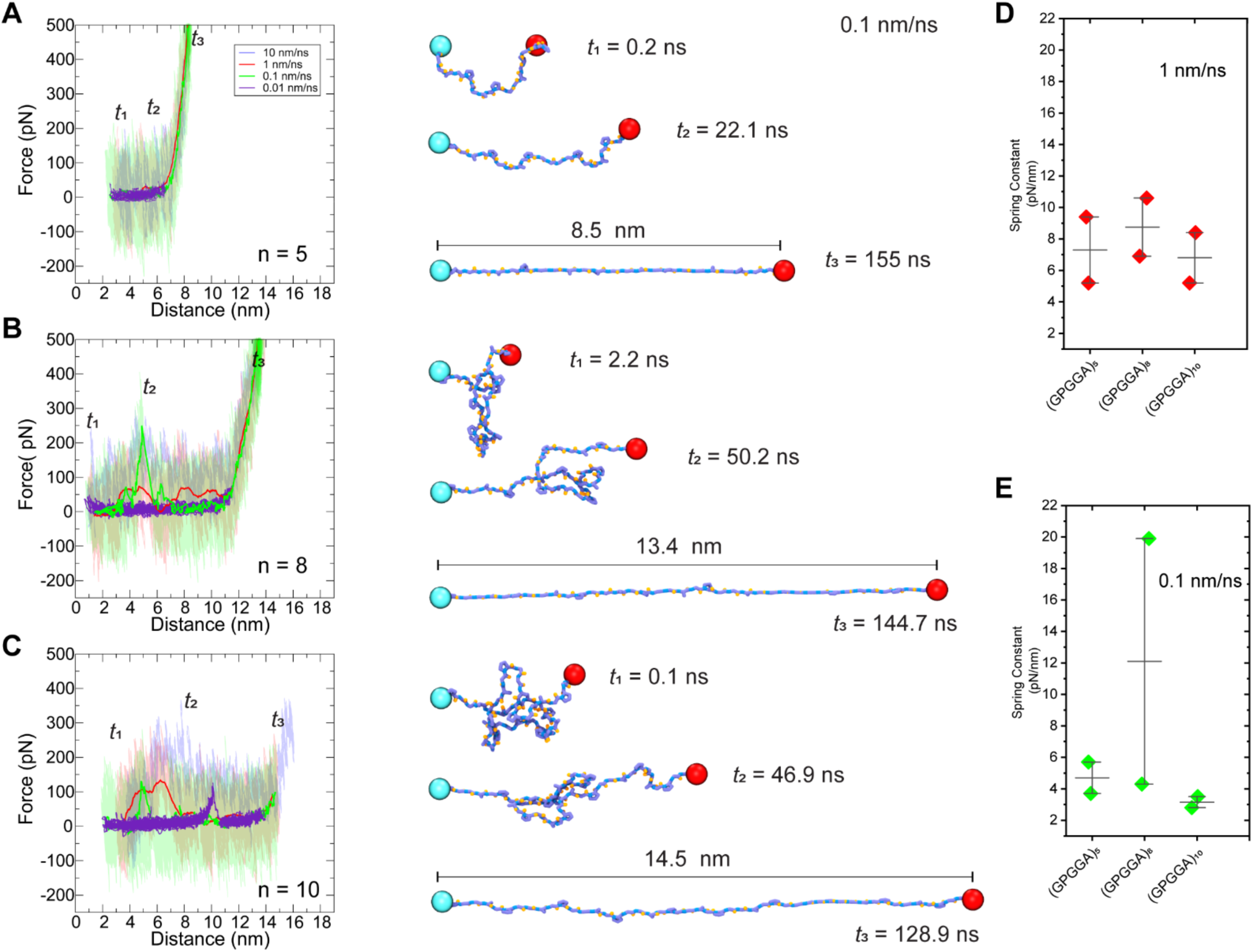
Elasticity of spider silk linkers (GPGGA)_n_ from constant-velocity SMD simulations. (*A*–*C*) Left panels show force-extension curves of (GPGGA)_n_ at different stretching velocities (0.01 nm/ns, 0.1 nm/ns, 1 nm/ns, and 10 nm/ns) for n = 5 (*A*), n = 7 (*B*), and n = 9 (*C*). These curves show two phases, with a first phase characterized by a gradual increase in force as the linkers stretch. In some cases, force peaks are observed. Linear regressions were used to obtain slope spring constants at 80% percent extension during the first phase (Fig. S3 and S4). Right panels show representative trajectory snapshots at indicated times as in Fig. 1 *B*. (*D*–*E*) Predicted spring constant values.

Similar elastic responses were observed for the longer spider silk peptides (GPGGA)_8_ and (GPGGA)_10_, with extensions that varied between ∼ 1 nm and ∼ 11 nm and between ∼ 2 nm and ∼ 14 nm, respectively (Fig. 2 *B-C* and S3 *B-C*). The simulation results and fits yielded similar spring constants to those predicted for the shorter version of the spider silk (Table 1). Interestingly, we did not observe a clear decrease in the value of spring constants with increasing length of the sensor (Figure 2 *D*-*E*), as theoretically expected and as experimentally measured (36). This might be caused by the lack of sequence information in the theoretical predictions. In addition, we observed the presence of force peaks, likely due to the formation of transient secondary structure (examined in detail subsequently), which were more pronounced at higher stretching velocities in our simulations. These force peaks were not observed in single molecule experiments in the low force region corresponding to straightening of the peptide (36), which is likely why our lower pulling rate simulations are more consistent with experimental results.

Force-extension curves for the synthetic linker systems (GGSGGS)_5_, (GGSGGS)_7_, and (GGSGGS)_9_ were similar to those predicted for the spider silk systems (Fig. 3, Table 1, and Figure S3 *D-F*). While some minor force peaks were still observed at higher pulling rates, these were less frequent and less prominent than for the spider silk sequence. Average values for spring constants were again comparable to theoretical values (Table 1), with trends that indicated a decrease in stiffness with decreased stretching velocity and decreased stiffness with increased length with exceptions, suggesting a tendency to deviate from a Hookean spring response to force, especially for the spider silk peptide.

**Figure 3.**
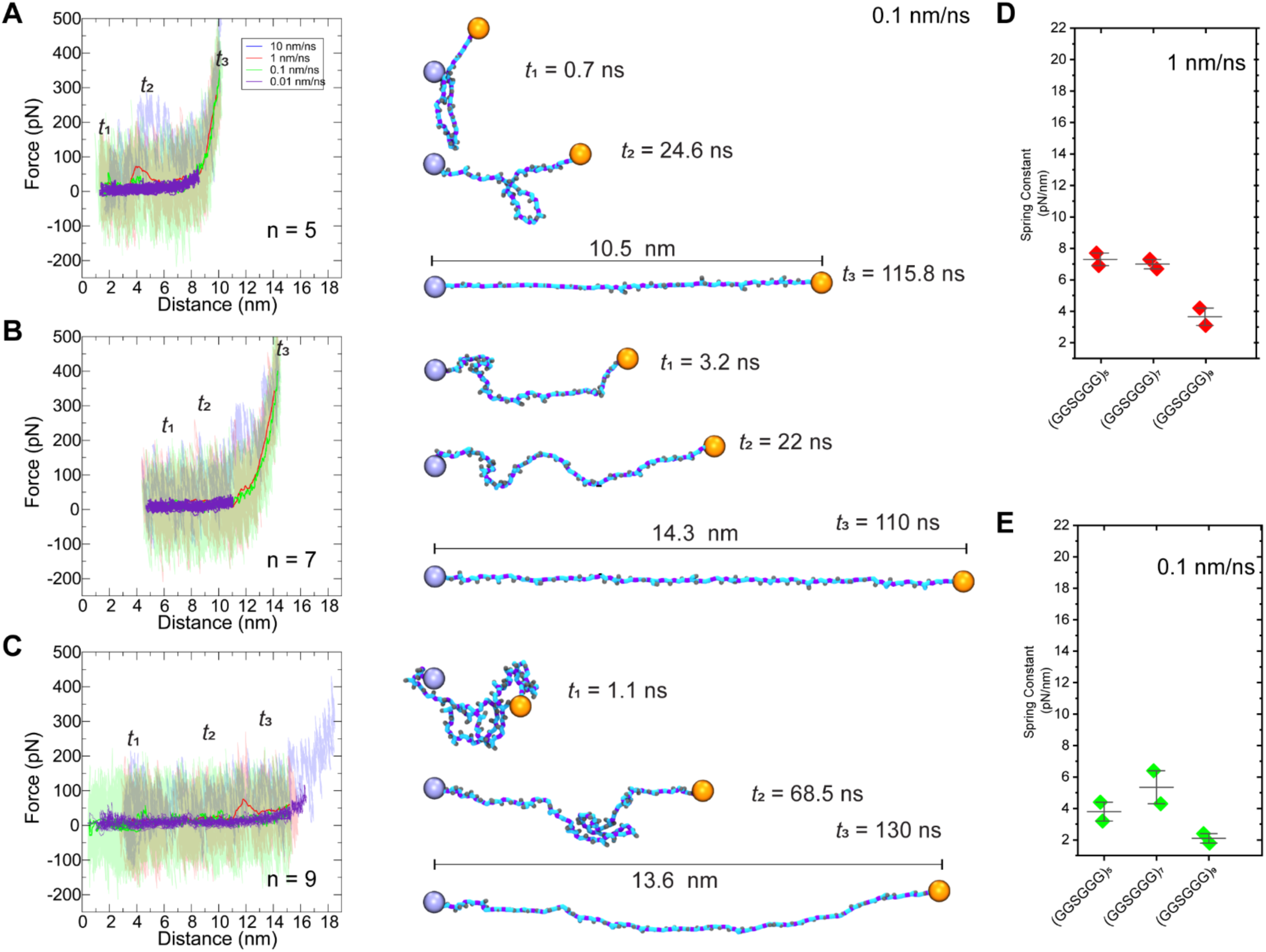
Elasticity of synthetic linkers (GGSGGS)_n_ from constant-velocity SMD simulations. (*A*–*C*) Left panels show force-extension curves of (GGSGGS)_n_ at different stretching velocities (0.01 nm/ns, 0.1 nm/ns, 1 nm/ns, and 10 nm/ns) for n = 5 (*A*), n = 7 (*B*), and n = 9 (*C*). These curves show two phases, with a first phase characterized by a gradual increase in force as the linkers stretch. Linear regressions were used to obtain slope spring constants at 80% percent extension during the first phase (Fig. S3 and S4). Right panels show representative trajectory snapshots at indicated times as in Fig. 1 *C*. (*D*–*E*) Predicted spring constant values.

We hypothesized that this deviation from theoretical predictions of the spider silk sequence was due to the formation of transient secondary structures during the extension that give rise to intermediate states that require force to disrupt. To further evaluate the source of force peaks, we carried out a comprehensive secondary structure analysis of the peptides during stretching simulations (Fig. 4, S5 and S6, Table S3, Video S4). We found that although peptide linkers predominantly remain unstructured, spider silk linkers exhibit transient extended ß hairpins approximately 4% of the time, in contrast to just 0.4% in synthetic linkers. This notable disparity likely explains the force peaks and higher slopes observed mainly in the force-distance curves of our SMD simulations for the spider-silk systems.

**Figure 4.**
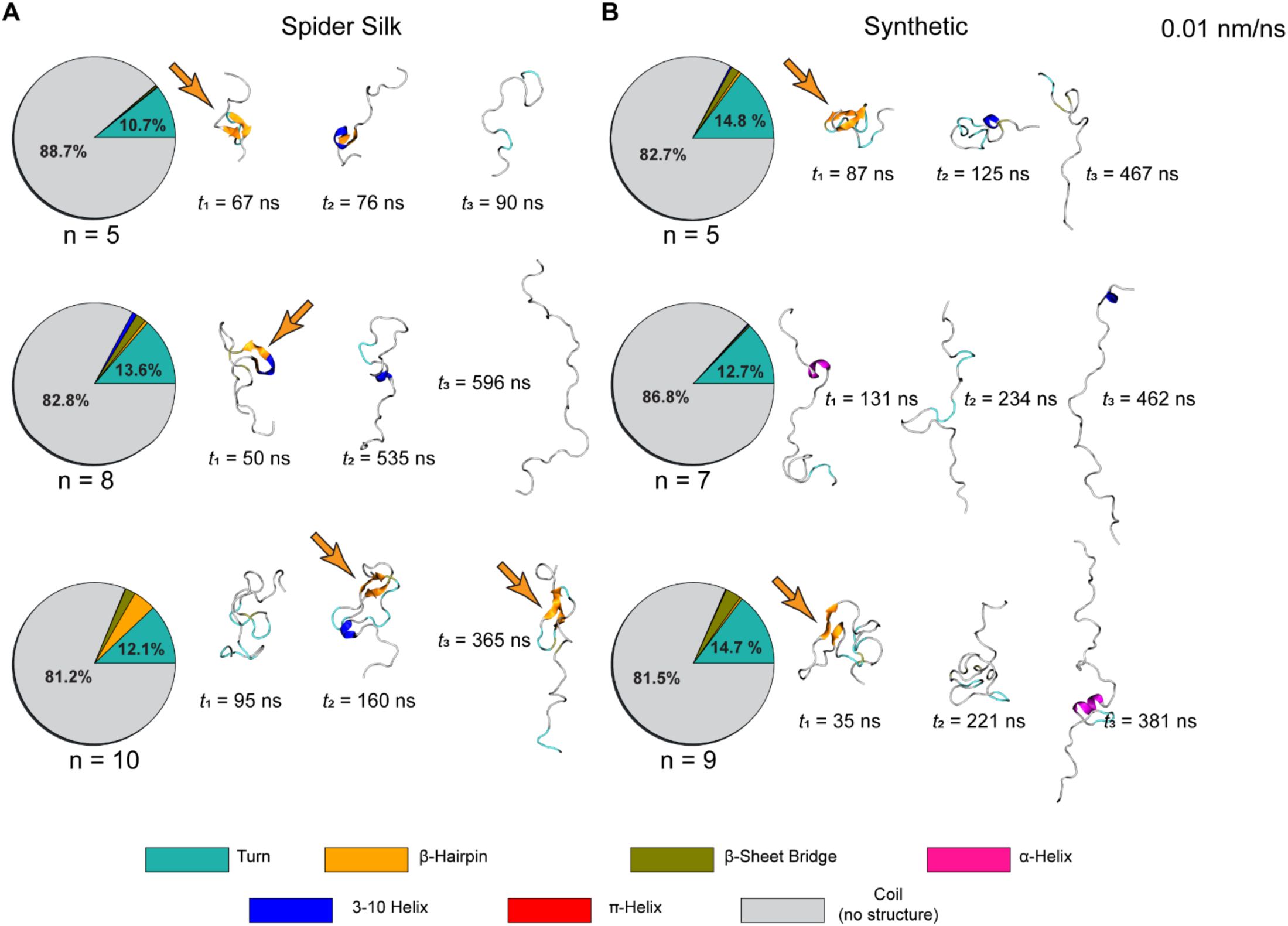
Secondary structure analysis of spider silk (GPGGA)_n_ and synthetic (GGSGGS)_n_ linkers during stretching. (*A-B*) Percentage of secondary structure motifs across all peptide-based linkers when stretched at 0.01 nm/ns. Pie charts showing percentage of secondary structure motifs across the entire trajectory for spider silk (GPGGA)_n_ (*A*) and synthetic (GGSGGS)_n_ peptides (*B*). Right panels show representative snapshots including secondary structure elements at indicated times.

### Elasticity of spider silk and synthetic linkers stretched at constant force

We also carried out constant-force simulations at 10 pN, 20 pN, 30 pN, and 50 pN for each of the six linker systems in duplicates (48 simulations; Tables S1-S2, Videos S5-S6). These simulations started from the same two initial conformations obtained during equilibrations of each system (10 ns and 15 ns) that served as starting points for constant-velocity SMD simulations. As expected, peptides extended from compact conformations to more extended states upon application of force (Fig. 5 and S7), although in some cases peptides fluctuated back to compact states at the lowest applied force (10 pN).

**Figure 5.**
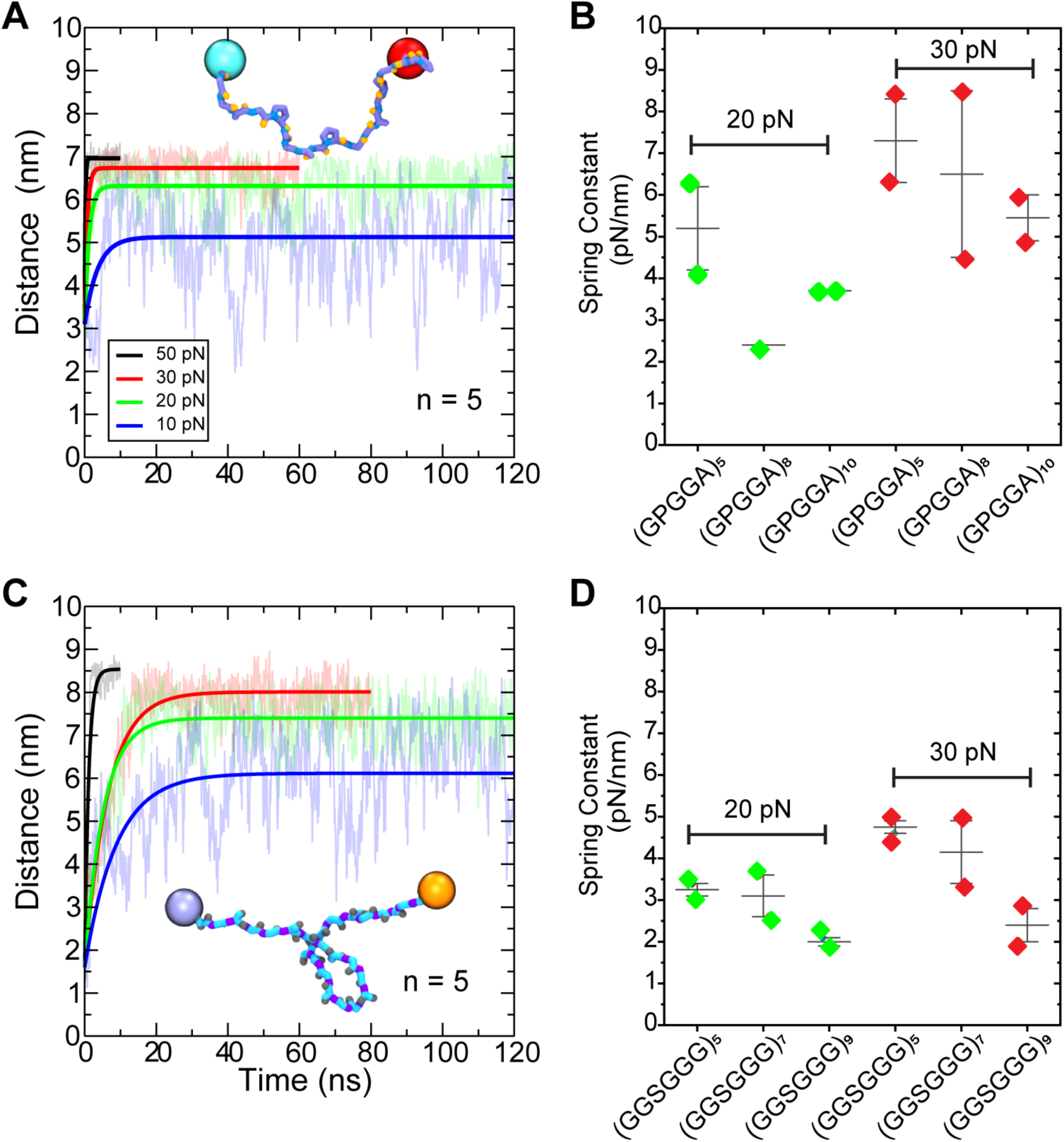
Elasticity of spider silk (GPGGA)_n_ and synthetic (GGSGGS)_n_ sensors from constant-force stretching. (*A*) Distance-time curves of SMD simulations performed at 10 pN (blue), 20 pN (green), 30 pN (red), and 50 pN (black) for (GPGGA)_5_ systems. Solid lines are fits used to obtain predicted spring constants. Insets show representative trajectory snapshots of each linker stretched at 20 pN as in Fig.1 *B*. (*B*) Spring constant values after pulling at a constant force of 20 pN and 30 pN for (GPGGA)_n_. (*C*) Distance-time curves of SMD simulations for (GGSGGS)_5_ shown as in *A.* (*D*) Spring constant values after pulling at a constant force of 20 pN and 30 pN for (GGSGGS)_n_ systems.

Distance-time curves for the smallest spider silk system (GPGGA)_5_ revealed rapid extensions of the linker at all forces tested, reaching final extensions that fluctuated around ∼ 5 nm, ∼ 6 nm, ∼ 7 nm, and ∼ 7 nm for forces of 10 pN 20 pN, 30 pN, and 50 pN, respectively (Fig. 5 *A* and S7 *A-C*). Interestingly, while the extension of this short peptide stretched at forces equal or larger than 20 pN fluctuated minimally around 6 nm to 7 nm after ∼ 10 ns, the extension at 10 pN fluctuated between ∼ 2 nm (compact state) and ∼ 6 nm (extended state) during the entire simulation. Similarly, distance-time curves for the longer spider silk systems (GPGGA)_8_ and (GPGGA)_10_ revealed rapid transitions from compact (∼ 3 nm for n = 8 and ∼ 4 nm for n = 10) to extended linker states (> 8 nm for n = 8 and > 12 nm for n = 10) at forces equal or larger than 20 pN (Fig. S7 *B-C*), but with transient intermediates at extensions of ∼ 5 nm (n = 8) and ∼ 7 nm (n = 10). When stretched at 10 pN, these longer peptides did not fully extend (n = 8) or transitioned between compact (∼ 4 nm), intermediate (∼ 6 nm), and fully extended (> 10 nm) states during the length of the simulations.

Estimates of the spring constants were obtained by fitting a mechanical model (see Methods) to the end-to-end distance versus time curves (Tables 2 and S1). In some instances, the presence of intermediates when using a stretching force of 10 pN prevented a clear fit. In particular, the longer constructs were dominated by more compact states at low force (Fig. 5 and S7). For the smaller system (GPGGA)_5_, spring constants increased with force as 3.7 ± 1.7 pN/nm, 5.2 ± 1.4 pN/nm, 7.3 ± 1.4 pN/nm, and 11.8 ± 1.8 pN/nm at 10, 20, 30, and 50 pN forces, respectively. These results reflect stiffening at larger forces as the backbone of the polypeptide chain starts to get stretched, explaining why values obtained at lower forces of 10 pN and 20 pN are more consistent with theoretical FJC predictions (4.03 pN/nm) and experimentally measured value (3.3 pN/nm) (36). Similarly, spring constant values for the longer spider silk systems (GPGGA)_8_ and (GPGGA)_10_ varied between 2.4 pN/nm and 10.5 pN/nm depending on stretching force and system. Interestingly, there was not clear decreasing trend in spring constant value as a function of peptide length when stretching all peptides at 20 pN and 30 pN (Fig. 5 *B*). likely due to secondary structure formation that occurs for the longer constructs.

Distance-time curves for the synthetic linker systems (GGSGGS)_5_, (GGSGGS)_7_, and (GGSGGS)_9_ were similar to those for the spider silk linker systems, with rapid extensions reaching values of ∼ 6 nm to ∼ 8 nm (n = 5), ∼ 8 nm to ∼ 12 nm (n = 7), and ∼12 nm to ∼ 16 nm (n = 9), at various forces from 10 pN to 50 pN (Fig. 5 *C* and S7). Transitions from compact states to extended and back were mainly observed for the longest linker (GGSGGS)_9_ stretched at 10 pN in both simulation repeats. Spring constant values for the synthetic linker systems ranged between 2 pN/m and 7.5 pN/m depending on length and stretching force (Tables 2, S2, S4). The simulation at 10 pN for the n = 9 system was not fitted because the linker exhibited large fluctuations between compact and extended states (Fig. S7). As with the spider silk linker systems, spring constant values for the synthetic systems increased with larger forces reflecting stretching of the polypeptide chain. Comparison of the spring constant values as a function of length for the synthetic systems did reveal a decreasing trend (Fig. 5 *D*), suggesting a Hookean spring response. In contrast, there is not a clear trend for the spider silk systems at any of the pulling forces and the (GPGGA)_8_ system showed the most variability.

Overall, constant-force simulations revealed that forces between 20 pN and 50 pN induced full linker stretching within the simulations’ time scales. In contrast, in simulations at 10 pN the linkers assumed various conformations without necessarily achieving full stretch after 240 ns, especially for the longer linkers (n = 7, 8, 9, 10). Spring constant values obtained from distance-time curves were mostly independent of initial conformation, with variations remaining minimal, specially at 20 pN and 30 pN for the synthetic linker systems, The spring constant values predicted from simulations (Table 2) remain comparable to theoretical and experimental values (Table 1) specially at pulling forces of 10 pN and 20 pN, likely because these lower forces do not cause significant backbone stretching. In general, the predicted spring constants from synthetic-peptide simulations exhibit less variability between starting conformations with an average standard deviation of 0.4 pN/nm in contrast to 1.7 pN/nm for the spider silk-peptide sensors, which is likely due to the presence of secondary structure in simulations of the spider silk peptide linkers.

**Table 2.**
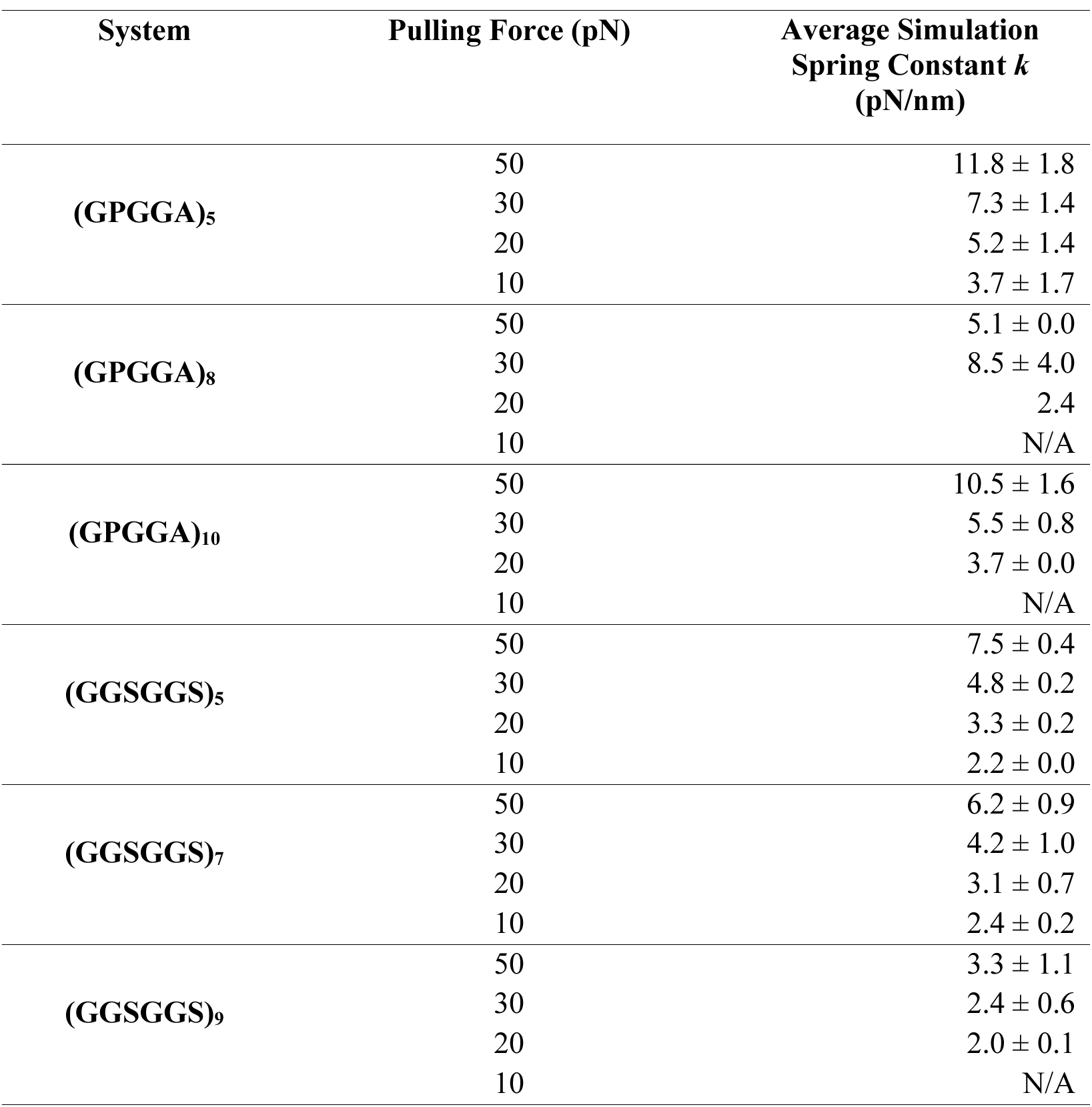
Average spring constants across all constant force simulations of (GPGGA)_n_ and (GGSGGS)_n_.

### Mechanical response of DNA sensors stretched at constant velocity and constant force

An all-atom model of an 18 bp dsDNA sensor (GGC CCG CAG CGA CCA CCC; Fig. 1 *D*) was equilibrated for 15 ns to allow the DNA duplex to relax into a native conformation. The resulting conformation was the starting point of all nine SMD simulations with constant-velocity stretching (Table S5). These simulations were done at 0.1 nm/ns, 1 nm/ns and 10 nm/ns by stretching in the positive *y* direction on the phosphorus atom of the terminal guanine of one strand and by stretching in the negative *y* direction on the complementary strand at three different locations: P1 (terminal cytosine), P2 (18^th^ guanine), and P3 (7^th^ guanine; Fig. 1 *E*). These three experimentally tested locations correspond to DNA unzipping (P1), DNA shearing (P2), and a mix between shearing and unzipping (P3) (22).

The force-extension curves for the DNA sensor stretched from P1 show small force peaks throughout the simulations, with similar force values at 0.1 nm/ns and 1 nm/ns and slightly larger forces at 10 nm/ns. These peaks correspond to DNA unzipping which requires a small force stimulus for full unfolding (Video S7) (45,46). The maximum force peaks were 163 pN, 191 pN, and 468 pN at stretching velocities of 0.1 nm/ns, 1 nm/ns, and 10 nm/ns respectively (Fig. 6 *A*).

**Figure 6.**
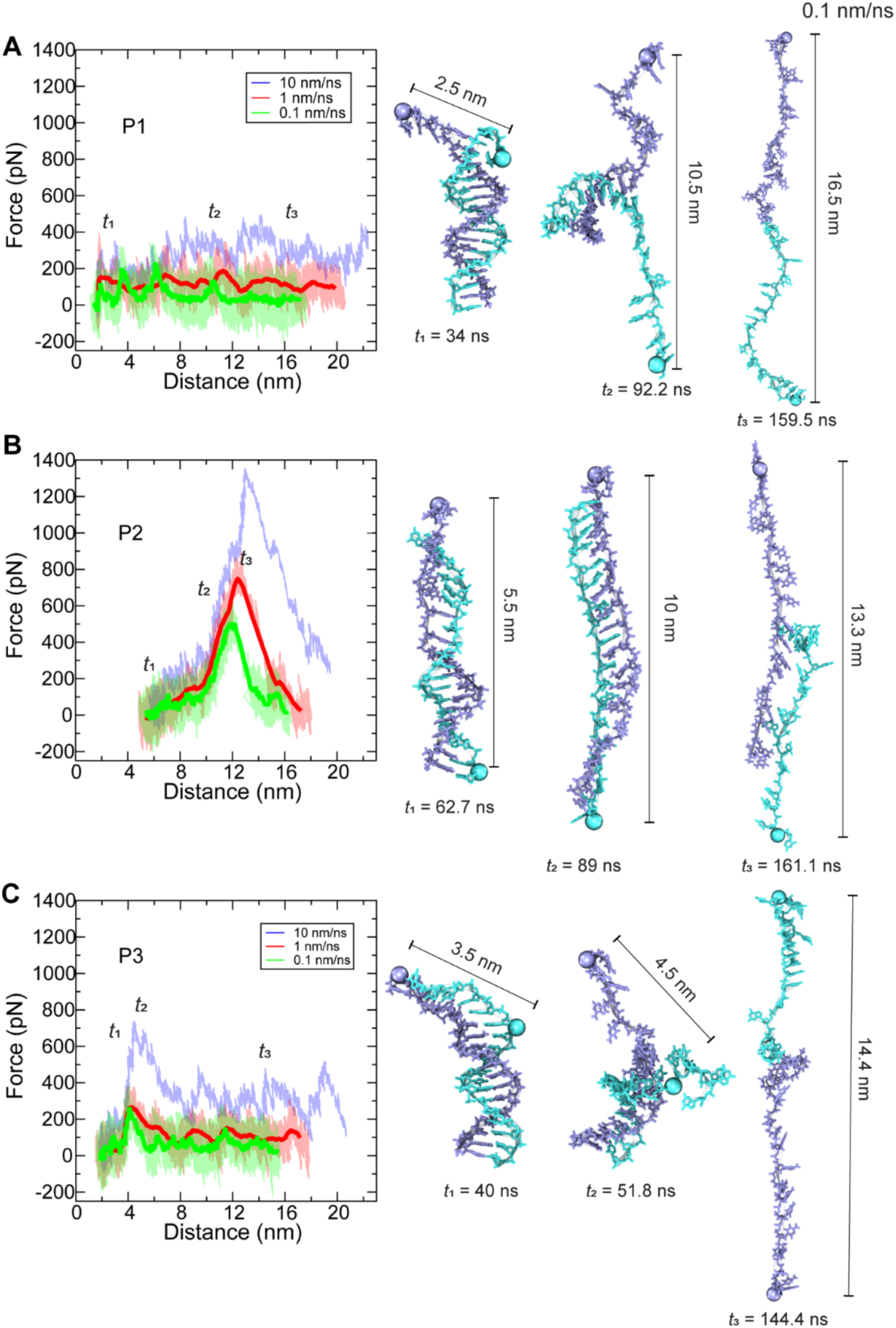
Mechanics of DNA-based sensors from constant-velocity SMD simulations. (*A*–*C*) Left panels show force-extension curves and 1-ns running averages (solid lines) of an 18-bp DNA force sensor stretched from different locations (P1, P2 and P3, respectively). At different stretching velocities (0.1 nm/ns, 1 nm/ns and 10 nm/ns). Right panels show representative trajectory snapshots at indicated times as in Fig. 1 *E*.

Unlike in simulations of the DNA sensor stretched from P1, stretching from P2 results in force-extension curves with two phases. In the first phase, the DNA sensor stretches from end-to-end distances of ∼ 4.5 nm to ∼8.5 nm with forces rising linearly from 0 pN to ∼100 pN, ∼160 pN, and ∼300 pN at different velocities (0.1 nm/ns, 1 nm/ns, and 10 nm/ns; Fig. 6 *B*). This phase corresponds to the elongation of the DNA duplex into a stretched state. In the second phase there is a sharp increase of forces leading to single peaks of varying magnitudes depending on stretching velocity with peak force values of 490 pN, 752 pN, and 1,242 pN for pulling rates of 0.1 nm/ns, 1 nm/ns, and 10 nm/ns, respectively (Fig. 6 *B*; Video S8).

Force-extension curves for the DNA sensor stretched from P3 exhibit features of both the unzipping (P1) and shearing (P2) modes. The initial force response resembles the shearing (P2) response with the presence of a major force peak at ∼4 to 4.5 nm end-to-end distances at all stretching velocities. The simulations reveal the DNA slightly rotates upon initial pulling and the large force peak corresponds to shearing of the first 7 bp between the two locations of pulling (Fig. 6 *C*, right). The DNA duplex then continues to dissociate base-pair by base-pair exhibiting unzipping behavior until it fully separates (Video S9). The maximum force peaks were 217 pN, 274 pN, and 652 pN for 0.1 nm/ns, 1 nm/ns, and 10 nm/ns stretching velocities, respectively (Fig. 6 *C*). Our simulations suggest that the maximum rupture force is governed by shearing of the short duplex region between the two pulling locations, and that unzipping would likely not play a role in any experimentally observed rupture. However, longer unzipping regions are likely important for the stability of the duplex.

We also did constant-force SMD simulations of the same DNA duplex using stretching locations P1, P2 and P3 and starting from the same conformations used for constant-velocity SMD simulations (12 simulations, Table S5). These simulations predict short-lived, transient intermediate states, with the unzipping configuration (P1) generally breaking faster than the shearing configuration (P2). The intermediate states are predicted to exist even at high forces expected to rupture these sensors (see Supplemental Discussion and Fig. S8), thus highlighting the relevance of observation timescales in interpreting experimental results.

Prior work has estimated the rupture forces of the dsDNA sensor to be in the range of ∼10-50 pN at near equilibrium conditions (22), with the unzipping configuration (P1) exhibiting lower rupture forces and the shearing configuration (P2) exhibiting higher rupture forces. Our constant-velocity simulations explored non-equilibrium conditions in which pulling rates are orders of magnitude larger than those typically used in single-molecule force spectroscopy and assumed in theoretical estimates of rupture force for DNA-based sensors. As predicted theoretically and computationally (47–52), and as experimentally verified using high-speed dynamic force spectroscopy with a high-speed AFM stretching proteins and protein-ligand systems (53,54), rupture force peak magnitudes increase with pulling rates, so forces at near-equilibrium conditions cannot be compared directly to forces from our constant-velocity SMD simulations. Similarly, transient events observed in constant-force simulations lasting hundreds of nanoseconds cannot be directly compared to events occurring over timescales of seconds. Nevertheless, both constant-velocity simulations at high loading rates and constant-force simulations using large forces during short timescales predict the same hierarchy of strengths expected at near equilibrium conditions, with the unzipping configuration (P1) exhibiting lower rupture forces and the shearing configuration (P2) exhibiting higher rupture forces. These predictions are applicable to cases in which loading rates are high and time scales are short, such as perception of loud high frequency sounds where stretching velocities of mechanosensitive proteins are expected to be as fast as 0.01 nm/ns and relevant physiological timescales can be on the order of tens of microseconds (55–57).

## DISCUSSION AND CONCLUSIONS

We explored the mechanical properties of three different types of molecular sensors—two peptide-based systems and one DNA-based. To this end, we used SMD simulations to stretch all systems at constant velocity and constant force. Simulation results showed that the peptide systems predicted stiffnesses are consistent with theoretical and experimental stiffnesses values, despite probing different force regimes (high loading rates, short timescales, and large forces for simulations versus near equilibrium conditions, small forces, and long timescales for theoretical and experimental estimates). Simulations for the DNA-based sensors predict larger rupture forces than those estimated at low loading rates, as expected. Interestingly, the hierarchy of strengths for different stretching configurations for DNA sensors is maintained, with DNA unzipping predicted to require less force than shearing also at the high loading rates used in our simulations.

We characterized mechanical properties of two common peptide-based sensors, the spider silk flagelliform-derived (GPGGA)_n_ and the synthetic (GGSGGS)_n_ sensors. Our findings indicate that the stiffnesses predicted from simulations and the expected theoretical values are in reasonable agreement. The presence of force fluctuations and force peaks in our fast constant-velocity stretching suggest a non-linear mechanical response that might be masked in quasi-equilibrium experiments. Constant-force simulations provide predictions that seem to be less variable. Regardless of the stretching mode, our data suggest that synthetic (GGSGGS)_n_ peptides exhibit more consistent linear responses, potentially making them preferable for force sensor designs. While both spider silk-based and synthetic peptide-based sensors have minimal secondary structure formation, the transient β- hairpin formed by the spider silk-based sensors appear to substantially influence their elasticity (Table S3; Video S4).

In addition to understanding the mechanical response of molecular force sensors and identifying simulation parameters that allow for reasonable comparison to experimental results, we also aim to provide insights that can guide the design of new force sensors. For example, our simulations suggest that the selection of fluorescent proteins can be optimized. Spring constant values for various peptides and selected FRET pair combinations vary significantly (Tables 3 and S7-S8) when considering distances where FRET efficiencies range between 20% and 80% (Table 2 and S9). The chosen FRET pair affects the distance range probed, which notably impacts stiffness estimates from constant-velocity simulations, making it a critical consideration in selecting linkers and fluorescent protein pairs for force sensing applications.

**Table 3.**
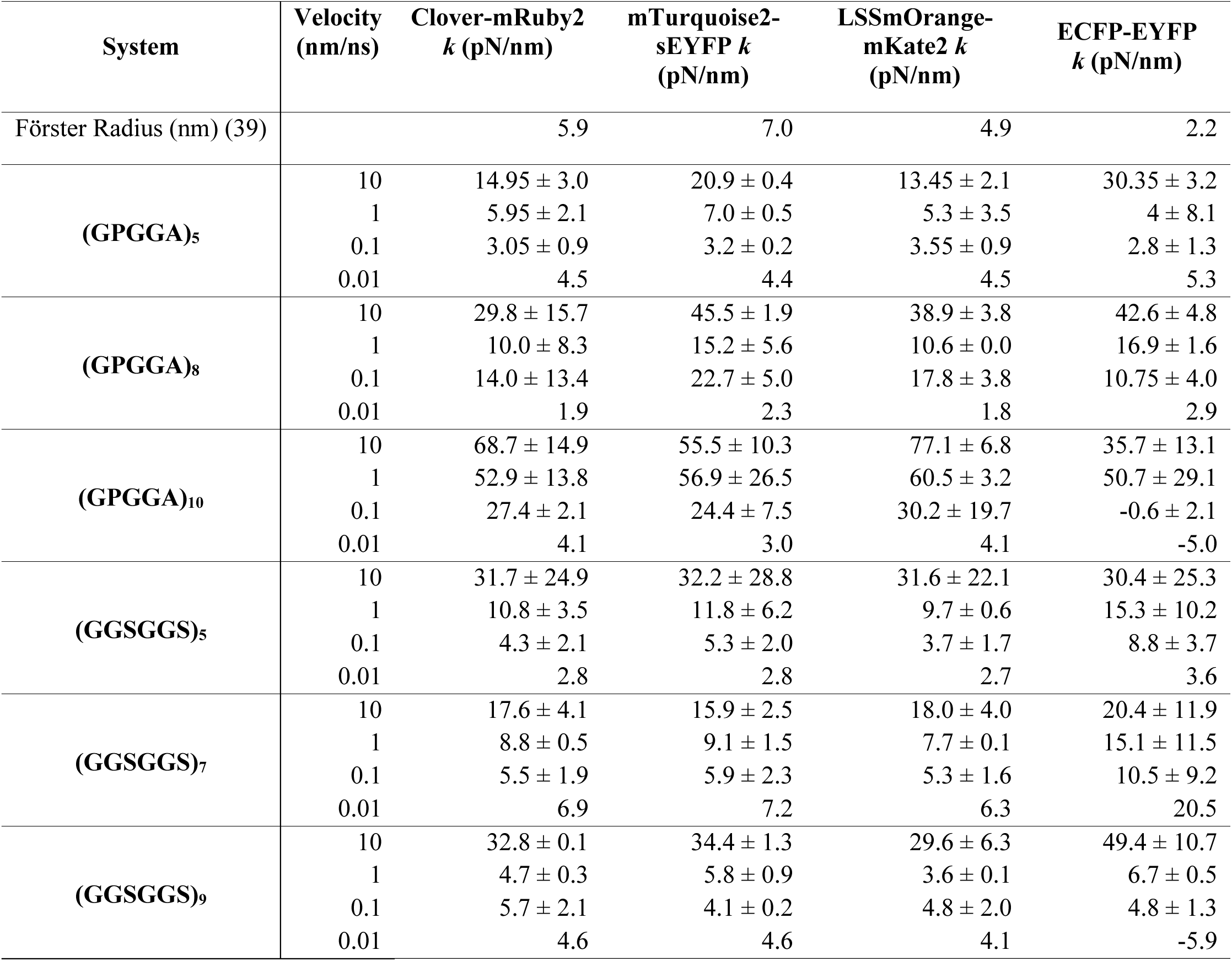
Average spring constants for 0.01 nm/ns constant-velocity simulations of (GPGGA)_n_ and (GGSGGS)_n_ at 80% extension and at relevant FRET distances.

| System | Velocity<br>(nm/ns) | Clover-mRuby2<br>*k* (pN/nm) | mTurquoise2-sEYFP *k*<br>(pN/nm) | LSSmOrange-<br>mKate2 *k*<br>(pN/nm) | ECFP-EYFP<br>*k* (pN/nm) |
| --- | --- | --- | --- | --- | --- |
| Förster Radius (nm) (39) |  | 5.9 | 7.0 | 4.9 | 2.2 |
| (GPGGA)<sub>5</sub> | 10 | 14.95 ± 3.0 | 20.9 ± 0.4 | 13.45 ± 2.1 | 30.35 ± 3.2 |
|  | 1 | 5.95 ± 2.1 | 7.0 ± 0.5 | 5.3 ± 3.5 | 4 ± 8.1 |
|  | 0.1 | 3.05 ± 0.9 | 3.2 ± 0.2 | 3.55 ± 0.9 | 2.8 ± 1.3 |
|  | 0.01 | 4.5 | 4.4 | 4.5 | 5.3 |
| (GPGGA)<sub>8</sub> | 10 | 29.8 ± 15.7 | 45.5 ± 1.9 | 38.9 ± 3.8 | 42.6 ± 4.8 |
|  | 1 | 10.0 ± 8.3 | 15.2 ± 5.6 | 10.6 ± 0.0 | 16.9 ± 1.6 |
|  | 0.1 | 14.0 ± 13.4 | 22.7 ± 5.0 | 17.8 ± 3.8 | 10.75 ± 4.0 |
|  | 0.01 | 1.9 | 2.3 | 1.8 | 2.9 |
| (GPGGA)<sub>10</sub> | 10 | 68.7 ± 14.9 | 55.5 ± 10.3 | 77.1 ± 6.8 | 35.7 ± 13.1 |
|  | 1 | 52.9 ± 13.8 | 56.9 ± 26.5 | 60.5 ± 3.2 | 50.7 ± 29.1 |
|  | 0.1 | 27.4 ± 2.1 | 24.4 ± 7.5 | 30.2 ± 19.7 | -0.6 ± 2.1 |
|  | 0.01 | 4.1 | 3.0 | 4.1 | -5.0 |
| (GGSGGS)<sub>5</sub> | 10 | 31.7 ± 24.9 | 32.2 ± 28.8 | 31.6 ± 22.1 | 30.4 ± 25.3 |
|  | 1 | 10.8 ± 3.5 | 11.8 ± 6.2 | 9.7 ± 0.6 | 15.3 ± 10.2 |
|  | 0.1 | 4.3 ± 2.1 | 5.3 ± 2.0 | 3.7 ± 1.7 | 8.8 ± 3.7 |
|  | 0.01 | 2.8 | 2.8 | 2.7 | 3.6 |
| (GGSGGS)<sub>7</sub> | 10 | 17.6 ± 4.1 | 15.9 ± 2.5 | 18.0 ± 4.0 | 20.4 ± 11.9 |
|  | 1 | 8.8 ± 0.5 | 9.1 ± 1.5 | 7.7 ± 0.1 | 15.1 ± 11.5 |
|  | 0.1 | 5.5 ± 1.9 | 5.9 ± 2.3 | 5.3 ± 1.6 | 10.5 ± 9.2 |
|  | 0.01 | 6.9 | 7.2 | 6.3 | 20.5 |
| (GGSGGS)<sub>9</sub> | 10 | 32.8 ± 0.1 | 34.4 ± 1.3 | 29.6 ± 6.3 | 49.4 ± 10.7 |
|  | 1 | 4.7 ± 0.3 | 5.8 ± 0.9 | 3.6 ± 0.1 | 6.7 ± 0.5 |
|  | 0.1 | 5.7 ± 2.1 | 4.1 ± 0.2 | 4.8 ± 2.0 | 4.8 ± 1.3 |
|  | 0.01 | 4.6 | 4.6 | 4.1 | -5.9 |

Our analyses of the mechanical properties of an 18 bp DNA sensor stretched from three locations confirm that DNA-based sensors are more suitable for detecting single force level corresponding to dissociation of the two strands rather than detecting a range of forces. Although the rupture force values we observe in our DNA sensor simulations are higher than experimentally observed, as expected because of the higher loading rate (47,49–54), the trends are consistent with experimental results (58), which report rupture forces that are low for unzipping (stretching forces P1) and are high for shearing (stretching from P2). Furthermore, the DNA sensor pulled from P3 is predicted to exhibit a mechanical response that is a combination of what we observed when stretching from P1 and P2 in simulations. In this case, the rupture force is dominated by shearing of the shorter part of the duplex region between the two pulling locations, with the DNA subdomain not directly stretched likely providing stability that allows the overall duplex to stay bound. We suggest that these types of simulations can be used to design and elucidate mechanical response of DNA-sensors of different sequences or lengths.

For instance, stretching of the DNA-sensor from P2 shows a Hookean spring-like behavior with extensions between 4.5 nm and 8.5 nm. This distance is relevant for the efficiency of many FRET pairs (e.g. cy5-cy3; Fig. S8). A linear fit on the extension region between 4.5 nm and 8.5 nm for the slowest stretching velocity (0.1 nm/ns) reveals a spring constant of ∼ 30 pN/nm (Fig. S9). This suggests that a dsDNA sensor with stretching locations as in P2 but with fluorophores placed between 2 nm to 4 nm apart, could provide a continuous or analog readout of force. However, further work would be required to explore whether the lack of stability and short lifetime of stretched DNA intermediates may preclude a practical realization of these type of sensors.

The velocities used in our SMD simulations (0.01 nm/ns – 10 nm/ns) surpass those used in some near equilibrium experiments but are close to those used in high-speed dynamic force spectroscopy using the high-speed AFM (up to 0.004 nm/ns) (54). Typically, fast stretching velocities in both computational and experimental force spectroscopy of proteins leads to large force peaks associated to rupture events (47–54,59). Consequently, it was unclear how much the pulling rate would affect a stiffness that is largely mediated by entropic elasticity of the coils in the case of the peptide-based sensors. We found that predicted stiffness values from simulations (Tables 1, S1 and S2) are in fair agreement with experimentally measured values. The lack of structure in these peptides-based sensors makes their predicted elasticity less susceptible to stretching velocity. This is in contrast to what we observed for DNA sensors, where rupture of hydrogen bonds seems to lead to force peak magnitudes that are, as expected, dependent on stretching velocities.

Our simulation results provide valuable insights for characterizing molecular sensors *in silico*. Constant-velocity SMD simulations of peptide sensors at 0.1 and 0.01 nm/ns predict elasticity and behavior that is less affected by stretching rates. These sensors might be ideal to explore both near equilibrium conditions and fast biological processes like those occurring in auditory mechanotransduction. DNA rupture force peaks are dependent on stretching velocities, and our simulations of DNA-based sensors at 0.1 nm/ns explore their response at high loading rates, not yet tested experimentally. Given that loading rates in physiological contexts are in many cases unknown (22), it is important to consider the mechanical response of sensors at both low and high stretching velocities and forces.

The simulations methods and results presented in here could also aid the design of new sensors with strategically placed FRET pairs, different sequences, or sequence lengths. Other applications and outlooks of this work include guiding the implementation of sensors in hybrid molecular devices requiring force sensitivity (58,60) or guiding tether design in force spectroscopy techniques like optical trapping or AFM, as demonstrated by Ott et al. (61).

## AUTHOR CONTRIBUTIONS

M.S., C.E.C., and D.M.L. designed the project. D.M.L. performed all simulations and data analyses. M.S. and C.E.C. supervised all work and provided support through the entire project. M.S. verified all the methods mentioned in this manuscript. D.M.L., M.S., and C.E.C wrote all sections of the manuscript. All authors discussed the results and contributed to the final manuscript.

## ACKNOWLEDGMENT

Simulations were performed at the Ohio Supercomputer Center Oakley supercomputers Owens and Pitzer (grants PAS1037 and PAA0217 to M.S.).

## ABBREVIATIONS

VMD: visual molecular dynamics
MD: molecular dynamics
AFM: atomic force microscopy
SMD: steered molecular dynamics
FRET: Förster resonance energy transfer

## SUPPORTING CITATIONS

References (39,62–64) appear in the Supporting material.

## REFERENCES

1. Bao, G. 2002. Mechanics of biomolecules. Journal of the Mechanics and Physics of Solids. 50(11):2237–2274, doi: 10.1016/s0022-5096(02)00035-2.

2. Pegoraro, A. F., P. Janmey, and D. A. Weitz. 2017. Mechanical Properties of the Cytoskeleton and Cells. Cold Spring Harbor Perspectives in Biology. 9(11):a022038, doi: 10.1101/cshperspect.a022038.

3. Egan, P., R. Sinko, P. R. Leduc, and S. Keten. 2015. The role of mechanics in biological and bio-inspired systems. Nature Communications. 6(1):7418, doi: 10.1038/ncomms8418.

4. Humphrey, J. D., and M. A. Schwartz. 2021. Vascular Mechanobiology: Homeostasis, Adaptation, and Disease. Annu Rev Biomed Eng. 23:1–27, doi: 10.1146/annurev-bioeng-092419-060810.

5. Kefauver, J. M., A. B. Ward, and A. Patapoutian. 2020. Discoveries in structure and physiology of mechanically activated ion channels. Nature. 587(7835):567–576, doi: 10.1038/s41586-020-2933-1.

6. Katta, S., M. Krieg, and M. B. Goodman. 2015. Feeling force: physical and physiological principles enabling sensory mechanotransduction. Annu Rev Cell Dev Biol. 31:347–371, doi: 10.1146/annurev-cellbio-100913-013426.

7. Vogel, V. 2006. Mechanotransduction involving multimodular proteins: converting force into biochemical signals. Annu Rev Biophys Biomol Struct. 35:459–488, doi: 10.1146/annurev.biophys.35.040405.102013.

8. Neuman, K. C., and A. Nagy. 2008. Single-molecule force spectroscopy: optical tweezers, magnetic tweezers and atomic force microscopy. Nat Methods. 5(6):491–505, doi: 10.1038/nmeth.1218, https://www.ncbi.nlm.nih.gov/pubmed/18511917.

9. Pimenta-Lopes, C., C. Suay-Corredera, D. Velázquez-Carreras, D. Sánchez-Ortiz, and J. Alegre-Cebollada. 2019. Concurrent atomic force spectroscopy. Communications Physics. 2(1), doi: 10.1038/s42005-019-0192-y.

10. Martucci, M., L. Debar, S. Van Den Wildenberg, and G. Farge. 2023. How to Quantify DNA Compaction by TFAM with Acoustic Force Spectroscopy and Total Internal Reflection Fluorescence Microscopy. In Springer US, pp. 121–137.

11. Woodside, M. T., P. C. Anthony, W. M. Behnke-Parks, K. Larizadeh, D. Herschlag, and S. M. Block. 2006. Direct measurement of the full, sequence-dependent folding landscape of a nucleic acid. Science. 314(5801):1001–1004, doi: 10.1126/science.1133601.

12. Mitgau, J., J. Franke, C. Schinner, G. Stephan, S. Berndt, D. G. Placantonakis, H. Kalwa, V. Spindler, C. Wilde, and I. Liebscher. 2022. The N Terminus of Adhesion G Protein-Coupled Receptor GPR126/ADGRG6 as Allosteric Force Integrator. Front Cell Dev Biol. 10:873278, doi: 10.3389/fcell.2022.873278.

13. Grashoff, C., B. D. Hoffman, M. D. Brenner, R. Zhou, M. Parsons, M. T. Yang, M. A. McLean, S. G. Sligar, C. S. Chen, T. Ha, and M. A. Schwartz. 2010. Measuring mechanical tension across vinculin reveals regulation of focal adhesion dynamics. Nature. 466(7303):263–266, doi: 10.1038/nature09198.

14. Blakely, B. L., C. E. Dumelin, B. Trappmann, L. M. McGregor, C. K. Choi, P. C. Anthony, V. K. Duesterberg, B. M. Baker, S. M. Block, D. R. Liu, and C. S. Chen. 2014. A DNA-based molecular probe for optically reporting cellular traction forces. Nat Methods. 11(12):1229–1232, doi: 10.1038/nmeth.3145.

15. Stabley, D. R., C. Jurchenko, S. S. Marshall, and K. S. Salaita. 2012. Visualizing mechanical tension across membrane receptors with a fluorescent sensor. Nature Methods. 9(1):64–67, doi: 10.1038/nmeth.1747.

16. Wang, X., and T. Ha. 2013. Defining Single Molecular Forces Required to Activate Integrin and Notch Signaling. Science. 340(6135):991–994, doi: 10.1126/science.1231041.

17. Morimatsu, M., A. H. Mekhdjian, A. S. Adhikari, and A. R. Dunn. 2013. Molecular tension sensors report forces generated by single integrin molecules in living cells. Nano Lett. 13(9):3985–3989, doi: 10.1021/nl4005145.

18. Zhang, Y., C. Ge, C. Zhu, and K. Salaita. 2014. DNA-based digital tension probes reveal integrin forces during early cell adhesion. Nature Communications. 5(1):5167, doi: 10.1038/ncomms6167.

19. Jurchenko, C., and K. S. Salaita. 2015. Lighting Up the Force: Investigating Mechanisms of Mechanotransduction Using Fluorescent Tension Probes. Molecular and Cellular Biology. 35(15):2570–2582, doi: 10.1128/MCB.00195-15.

20. Hu, Y., Y. Duan, and K. Salaita. 2023. DNA Nanotechnology for Investigating Mechanical Signaling in the Immune System. Angewandte Chemie International Edition. 62(30):e202302967, doi: 10.1002/anie.202302967, https://onlinelibrary.wiley.com/doi/abs/10.1002/anie.202302967.

21. Morimatsu, M., A. H. Mekhdjian, A. S. Adhikari, and A. R. Dunn. 2013. Molecular Tension Sensors Report Forces Generated by Single Integrin Molecules in Living Cells. Nano Letters. 13(9):3985–3989, doi: 10.1021/nl4005145.

22. Wang, X., Z. Rahil, I. T. S. Li, F. Chowdhury, D. E. Leckband, Y. R. Chemla, and T. Ha. 2016. Constructing modular and universal single molecule tension sensor using protein G to study mechano-sensitive receptors. Scientific Reports. 6(1):21584, doi: 10.1038/srep21584.

23. Lacroix, A. S., A. D. Lynch, M. E. Berginski, and B. D. Hoffman. 2018. Tunable molecular tension sensors reveal extension-based control of vinculin loading. eLife. 7, doi: 10.7554/elife.33927.

24. Huang, Y., T. Chen, X. Chen, X. Chen, J. Zhang, S. Liu, M. Lu, C. Chen, X. Ding, C. Yang, R. Huang, and Y. Song. 2024. Decoding Biomechanical Cues Based on DNA Sensors. Small. doi: 10.1002/smll.202310330.

25. Heim, M., D. Keerl, and T. Scheibel. 2009. Spider silk: from soluble protein to extraordinary fiber. Angew Chem Int Ed Engl. 48(20):3584–3596, doi: 10.1002/anie.200803341.

26. Widhe, M., H. Bysell, S. Nystedt, I. Schenning, M. Malmsten, J. Johansson, A. Rising, and M. Hedhammar. 2010. Recombinant spider silk as matrices for cell culture. Biomaterials. 31(36):9575–9585, doi: 10.1016/j.biomaterials.2010.08.061.

27. Malay, A. D., H. C. Craig, J. Chen, N. A. Oktaviani, and K. Numata. 2022. Complexity of Spider Dragline Silk. Biomacromolecules. 23(5):1827–1840, doi: 10.1021/acs.biomac.1c01682.

28. Oroudjev, E., J. Soares, S. Arcidiacono, J. B. Thompson, S. A. Fossey, and H. G. Hansma. 2002. Segmented nanofibers of spider dragline silk: Atomic force microscopy and single-molecule force spectroscopy. Proceedings of the National Academy of Sciences. 99(suppl_2):6460–6465, doi: doi:10.1073/pnas.082526499, https://www.pnas.org/doi/abs/10.1073/pnas.082526499.

29. Pacios, L. F., J. Arguelles, C. Y. Hayashi, G. V. Guinea, M. Elices, and J. Perez-Rigueiro. 2022. Differences in the Elastomeric Behavior of Polyglycine-Rich Regions of Spidroin 1 and 2 Proteins. Polymers. 14(23):5263, https://www.mdpi.com/2073-4360/14/23/5263.

30. C. Bittencourt, D. M., P. F. Oliveira, B. M. Souto, S. M. Freitas, L. P. Silva, A. M. Murad, V. A. Michalczechen-Lacerda, R. V. Lewis, and E. L. Rech. 2021. Molecular Dynamics of Synthetic Flagelliform Silk Fiber Assembly. Macromolecular Materials and Engineering. 306(1):2000530, doi: 10.1002/mame.202000530.

31. Evers, T. H., E. M. W. M. Van Dongen, A. C. Faesen, E. W. Meijer, and M. Merkx. 2006. Quantitative Understanding of the Energy Transfer between Fluorescent Proteins Connected via Flexible Peptide Linkers. Biochemistry. 45(44):13183–13192, doi: 10.1021/bi061288t.

32. Mosayebi, M., A. A. Louis, J. P. K. Doye, and T. E. Ouldridge. 2015. Force-Induced Rupture of a DNA Duplex: From Fundamentals to Force Sensors. ACS Nano. 9(12):11993–12003, doi: 10.1021/acsnano.5b04726.

33. Mishra, R. K., and L. Maganti. 2022. Antitumor drugs effect on the stability of double-stranded DNA: steered molecular dynamics analysis. Journal of Biomolecular Structure and Dynamics. 40(21):11373–11382, doi: 10.1080/07391102.2021.1960193.

34. Raghunathan, K., J. N. Milstein, and J.-C. Meiners. 2011. Stretching Short Sequences of DNA with Constant Force Axial Optical Tweezers. Journal of Visualized Experiments.(56), doi: 10.3791/3405, http://europepmc.org/articles/pmc3227192?pdf=render.

35. Karplus, M., and G. A. Petsko. 1990. Molecular dynamics simulations in biology. Nature. 347(6294):631–639, doi: 10.1038/347631a0.

36. Brenner, M. D., R. Zhou, D. E. Conway, L. Lanzano, E. Gratton, M. A. Schwartz, and T. Ha. 2016. Spider Silk Peptide Is a Compact, Linear Nanospring Ideal for Intracellular Tension Sensing. Nano Letters. 16(3):2096–2102, doi: 10.1021/acs.nanolett.6b00305.

37. Ringer, P., A. Weißl, A.-L. Cost, A. Freikamp, B. Sabass, A. Mehlich, M. Tramier, M. Rief, and C. Grashoff. 2017. Multiplexing molecular tension sensors reveals piconewton force gradient across talin-1. Nature Methods. 14(11):1090–1096, doi: 10.1038/nmeth.4431.

38. Isralewitz, B., J. Baudry, J. Gullingsrud, D. Kosztin, and K. Schulten. 2001. Steered molecular dynamics investigations of protein function. Journal of Molecular Graphics and Modelling. 19(1):13–25, doi: 10.1016/s1093-3263(00)00133-9.

39. Bajar, B. T., E. S. Wang, S. Zhang, M. Z. Lin, and J. Chu. 2016. A Guide to Fluorescent Protein FRET Pairs. Sensors (Basel). 16(9), doi: 10.3390/s16091488.

40. Emsley, P., B. Lohkamp, W. G. Scott, and K. Cowtan. 2010. Features and development ofCoot. Acta Crystallographica Section D Biological Crystallography. 66(4):486–501, doi: 10.1107/s0907444910007493.

41. Munteanu, M. G., K. Vlahovicek, S. Parthasarathy, I. Simon, and S. Pongor. 1998. Rod models of DNA: sequence-dependent anisotropic elastic modelling of local bending phenomena. Trends Biochem Sci. 23(9):341–347, doi: 10.1016/s0968-0004(98)01265-1.

42. Humphrey, W., A. Dalke, and K. Schulten. 1996. VMD: Visual molecular dynamics. Journal of Molecular Graphics. 14(1):33–38, doi: 10.1016/0263-7855(96)00018-5.

43. Phillips, J. C., R. Braun, W. Wang, J. Gumbart, E. Tajkhorshid, E. Villa, C. Chipot, R. D. Skeel, L. Kalé, and K. Schulten. 2005. Scalable molecular dynamics with NAMD. Journal of Computational Chemistry. 26(16):1781–1802, doi: 10.1002/jcc.20289, https://onlinelibrary.wiley.com/doi/abs/10.1002/jcc.20289.

44. Huang, J., and A. D. Mackerell. 2013. CHARMM36 all-atom additive protein force field: Validation based on comparison to NMR data. Journal of Computational Chemistry. 34(25):2135–2145, doi: 10.1002/jcc.23354, http://europepmc.org/articles/pmc3800559?pdf=render.

45. Zhang, J., Y. Yan, S. Samai, and D. S. Ginger. 2016. Dynamic Melting Properties of Photoswitch-Modified DNA: Shearing versus Unzipping. The Journal of Physical Chemistry B. 120(41):10706–10713, doi: 10.1021/acs.jpcb.6b08297.

46. Tee, S. R., and Z. Wang. 2018. How Well Can DNA Rupture DNA? Shearing and Unzipping Forces inside DNA Nanostructures. ACS Omega. 3(1):292–301, doi: 10.1021/acsomega.7b01692.

47. Evans, E., and K. Ritchie. 1997. Dynamic strength of molecular adhesion bonds. Biophys J. 72(4):1541–1555, doi: 10.1016/s0006-3495(97)78802-7.

48. Bell, G. I. 1978. Models for the specific adhesion of cells to cells. Science. 200(4342):618–627, doi: 10.1126/science.347575.

49. Evans, E. A., and D. A. Calderwood. 2007. Forces and bond dynamics in cell adhesion. Science. 316(5828):1148–1153, doi: 10.1126/science.1137592.

50. Sotomayor, M., and K. Schulten. 2007. Single-molecule experiments in vitro and in silico. Science. 316(5828):1144–1148, doi: 10.1126/science.1137591.

51. Lee, E. H., J. Hsin, M. Sotomayor, G. Comellas, and K. Schulten. 2009. Discovery through the computational microscope. Structure. 17(10):1295–1306, doi: 10.1016/j.str.2009.09.001.

52. Dudko, O. K., G. Hummer, and A. Szabo. 2008. Theory, analysis, and interpretation of single-molecule force spectroscopy experiments. Proc Natl Acad Sci U S A. 105(41):15755–15760, doi: 10.1073/pnas.0806085105.

53. Rico, F., L. Gonzalez, I. Casuso, M. Puig-Vidal, and S. Scheuring. 2013. High-speed force spectroscopy unfolds titin at the velocity of molecular dynamics simulations. Science. 342(6159):741–743, doi: 10.1126/science.1239764.

54. Rico, F., A. Russek, L. González, H. Grubmüller, and S. Scheuring. 2019. Heterogeneous and rate-dependent streptavidin-biotin unbinding revealed by high-speed force spectroscopy and atomistic simulations. Proc Natl Acad Sci U S A. 116(14):6594–6601, doi: 10.1073/pnas.1816909116.

55. Corey, D. P., and A. J. Hudspeth. 1983. Kinetics of the receptor current in bullfrog saccular hair cells. J Neurosci. 3(5):962–976, doi: 10.1523/jneurosci.03-05-00962.1983.

56. Robles, L., and M. A. Ruggero. 2001. Mechanics of the mammalian cochlea. Physiol Rev. 81(3):1305–1352, doi: 10.1152/physrev.2001.81.3.1305.

57. Jaiganesh, A., Y. Narui, R. Araya-Secchi, and M. Sotomayor. 2018. Beyond Cell-Cell Adhesion: Sensational Cadherins for Hearing and Balance. Cold Spring Harb Perspect Biol. 10(9), doi: 10.1101/cshperspect.a029280.

58. Kurus, N. N., and F. N. Dultsev. 2018. Determination of the Thermodynamic Parameters of DNA Double Helix Unwinding with the Help of Mechanical Methods. ACS Omega. 3(3):2793–2797, doi: 10.1021/acsomega.7b01815.

59. Sheridan, S., F. Gräter, and C. Daday. 2019. How Fast Is Too Fast in Force-Probe Molecular Dynamics Simulations? The Journal of Physical Chemistry B. 123(17):3658–3664, doi: 10.1021/acs.jpcb.9b01251.

60. Le, J. V., Y. Luo, M. A. Darcy, C. R. Lucas, M. F. Goodwin, M. G. Poirier, and C. E. Castro. 2016. Probing Nucleosome Stability with a DNA Origami Nanocaliper. ACS Nano. 10(7):7073–7084, doi: 10.1021/acsnano.6b03218, https://www.ncbi.nlm.nih.gov/pubmed/27362329.

61. Ott, W., M. A. Jobst, M. S. Bauer, E. Durner, L. F. Milles, M. A. Nash, and H. E. Gaub. 2017. Elastin-like Polypeptide Linkers for Single-Molecule Force Spectroscopy. ACS Nano. 11(6):6346–6354, doi: 10.1021/acsnano.7b02694.

62. Danilowicz, C., V. W. Coljee, C. Bouzigues, D. K. Lubensky, D. R. Nelson, and M. Prentiss. 2003. DNA unzipped under a constant force exhibits multiple metastable intermediates. Proceedings of the National Academy of Sciences. 100(4):1694–1699, doi: 10.1073/pnas.262789199.

63. Bajar, B. T., E. S. Wang, A. J. Lam, B. B. Kim, C. L. Jacobs, E. S. Howe, M. W. Davidson, M. Z. Lin, and J. Chu. 2016. Improving brightness and photostability of green and red fluorescent proteins for live cell imaging and FRET reporting. Scientific Reports. 6(1):20889, doi: 10.1038/srep20889.

64. Son, H., W. Mo, J. Park, J.-W. Lee, and S. Lee. 2020. Single-Molecule FRET Detection of Sub-Nanometer Distance Changes in the Range below a 3-Nanometer Scale. Biosensors. 10(11):168, https://www.mdpi.com/2079-6374/10/11/168.

