## Supplementary text, figures, and tables for "*In-Silico* Analyses of Molecular Force Sensors for Mechanical Characterization of Biological Systems"

**Video S1** Equilibration of spider silk peptide. The $(GPGGA)_{10}$ peptide starts in an extended conformation and rapidly adopts a compact configuration. Peptide is shown in sticks with carbon atoms in gray, nitrogen atoms in brown, and oxygens in tan. Water molecules are omitted for clarity.

**Video S2** Stretching of spider silk $(GPGGA)_{10}$ peptide at constant velocity (0.1 nm/ns). Shown as in Video S1.

**Video S3** Stretching of synthetic $(GGSGGS)_{9}$ peptide at constant velocity (0.1 nm/ns). Shown as in Video S1.

**Video S4** Stretching of spider silk peptide at constant velocity highlighting secondary structure representations.

**Video S5** Stretching of spider silk $(GPGGA)_{10}$ peptide at constant force (20 pN). Shown as in Video S1.

**Video S6** Stretching of synthetic peptide $(GGSGGS)_{9}$ at constant force (20 pN). Shown as in Video S1.

**Video S7** DNA sensor stretching from P1 at constant velocity (0.1 nm/ns). The DNA duplex unzips. DNA shown in cartoon.

**Video S8** DNA sensor stretching from P2 at constant velocity (0.1 nm/ns). The DNA duplex slightly rotates, then elongates before the rupture of all base pairs simultaneously (shearing). DNA shown in cartoon and sticks.

**Video S9** DNA sensor stretching from P3 at constant velocity (0.1 nm/ns). The DNA duplex slightly rotates before multiple base pairs rupture simultaneously, then continues unzipping as shown as in Video S7.

**Mechanical response of DNA sensors stretched at constant force.**

The extension-time curves of the DNA sensor stretched from P1 (unzipping) show the presence of transient intermediates when using stretching forces of 50 pN and 100 pN, these intermediates remain for a longer time when stretched with a force of 50 pN. Unlike in the stretching conditions P2 (shearing) and P3 (mixed), the DNA sensor stretched from P1 fully ruptures at all stretching forces (50 pN 100 pN, 300 pN, and 500 pN; Fig. S8 *A*). The DNA sensor stretched from P2 only reaches full rupture at stretching forces of 300 pN and 500 pN (Fig. S8 *B*). The extension-time curves of the DNA sensor stretched from P2 exhibit intermediates at stretching forces of 300 pN, 100 pN and 50 pN. All intermediates are present for significantly longer times in comparison to P1 and P3 stretching conditions.

Lastly, in the extension-time curves of the DNA sensor stretched from P3 (mixed) we also observe the presence of intermediates at stretching forces of 100 pN and 50 pN, however the intermediates are present for longer times at both stretching forces (50 pN and 100 pN) in comparison to the stretching condition P1. The DNA sensor stretched from P3 reaches full rupture at all stretching forces except for 50 pN (Fig. S8 *C*). This behavior is in agreement with the experimental findings by Danilowicz (62).

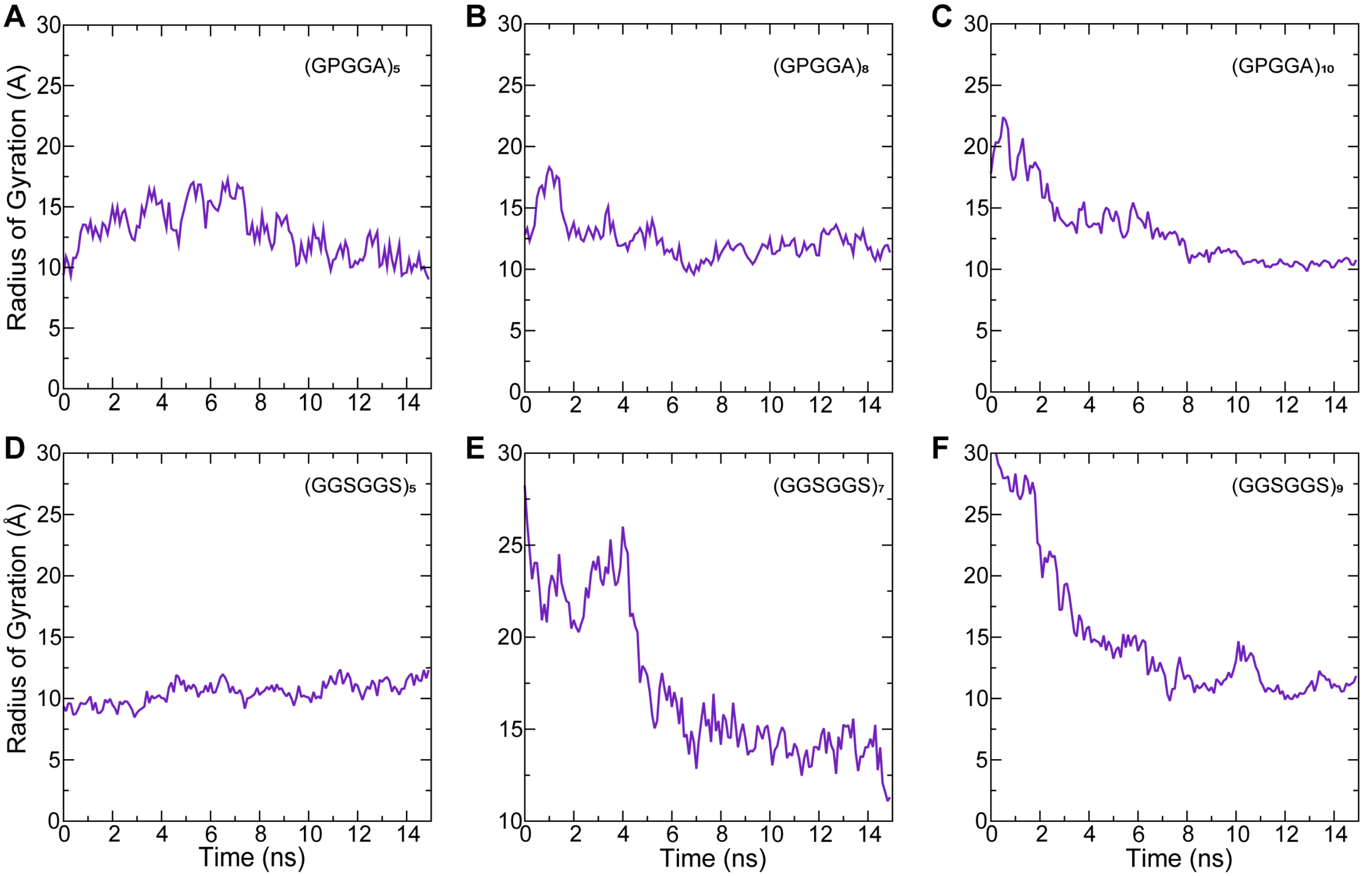


**Figure S1. Radius of gyration for peptides-based linkers during equilibration.** (*A–F*) Radius of gyration as a function of time for (GPGGA)$_n$ (*A–C*) and (GGSGGS)$_n$ (*B–F*) linkers.

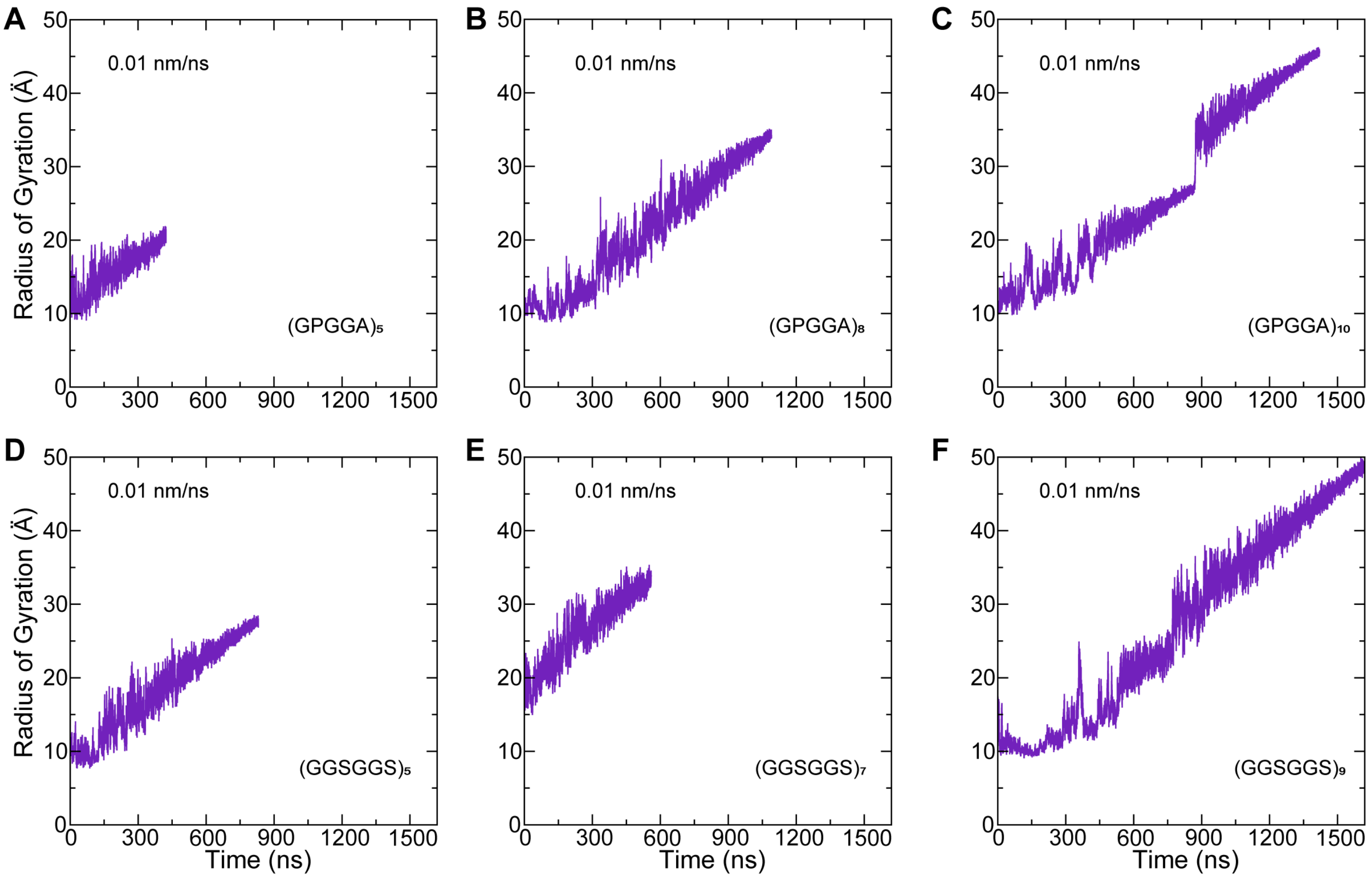


**Figure S2. Radius of gyration of (GPGGA)$_n$ and (GGSGGS)$_n$ peptides.** Radius of gyration as a function of time during 0.01 nm/ns constant-velocity stretching simulations. (*A-F*) Radius of gyration increases linearly as the peptides stretch. Some fluctuations can be observed in panels C, E, and F and are likely due to the formation of secondary structure.

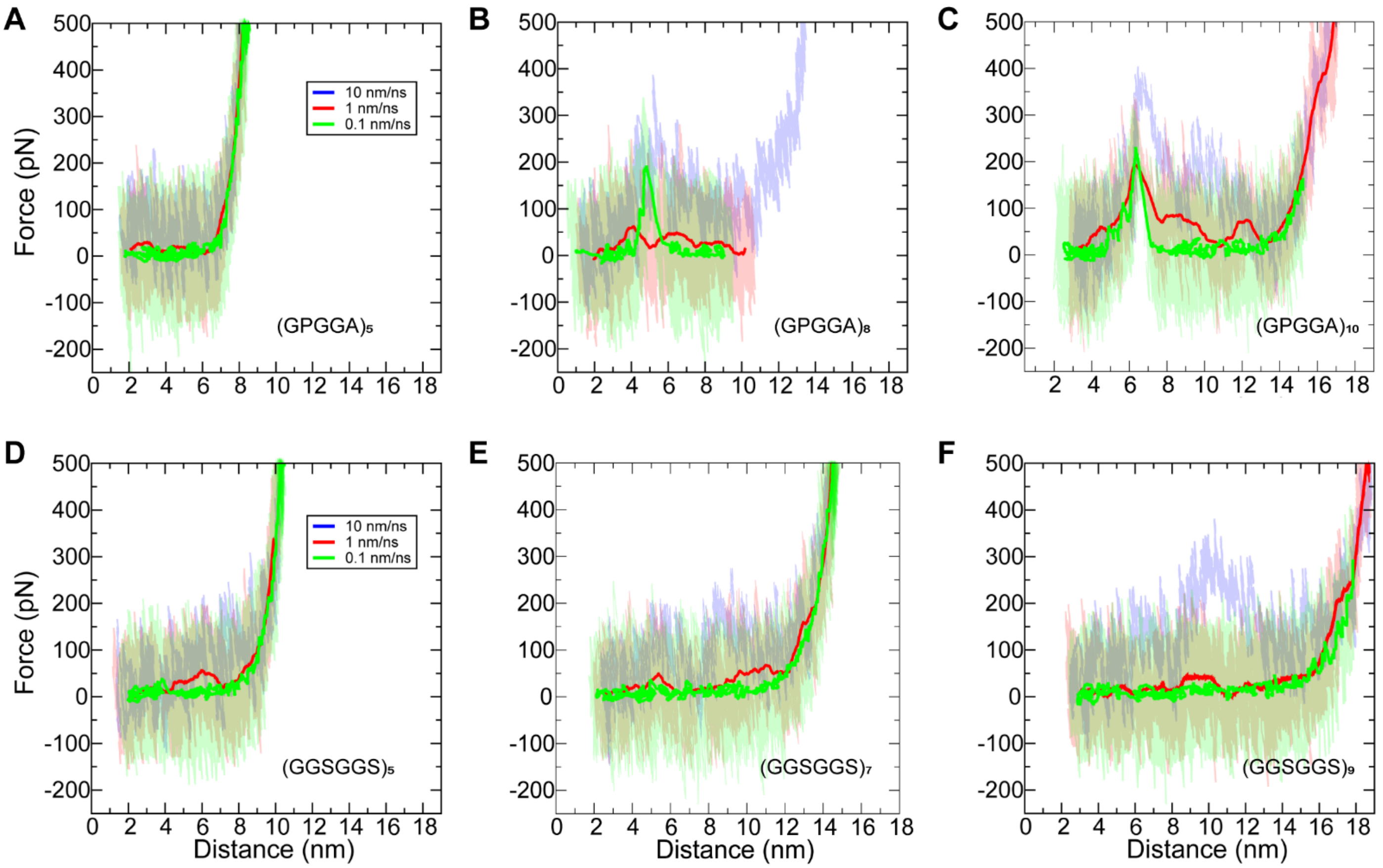


**Figure S3. Elasticity profiles of (GPGGA)$_n$ and (GGSGGS)$_n$ obtained from constant-velocity SMD simulations after 15 ns equilibration.** (*A-F*) Force-extension curves of peptide-based linkers stretched at different velocities (0.1 nm/ns, 1 nm/ns, and 10 nm/ns) for (GPGGA)$_5$ (*A*)*,* (GPGGA)$_8$ (*B*), (GPGGA)$_{10}$ (*C*), (GGSGGS)$_5$ (*D*), (GGSGGS)$_7$ (*E*), (GGSGGS)$_9$ (*F*). Extension curves in (*B*-*C*) show force peaks across each trajectory.

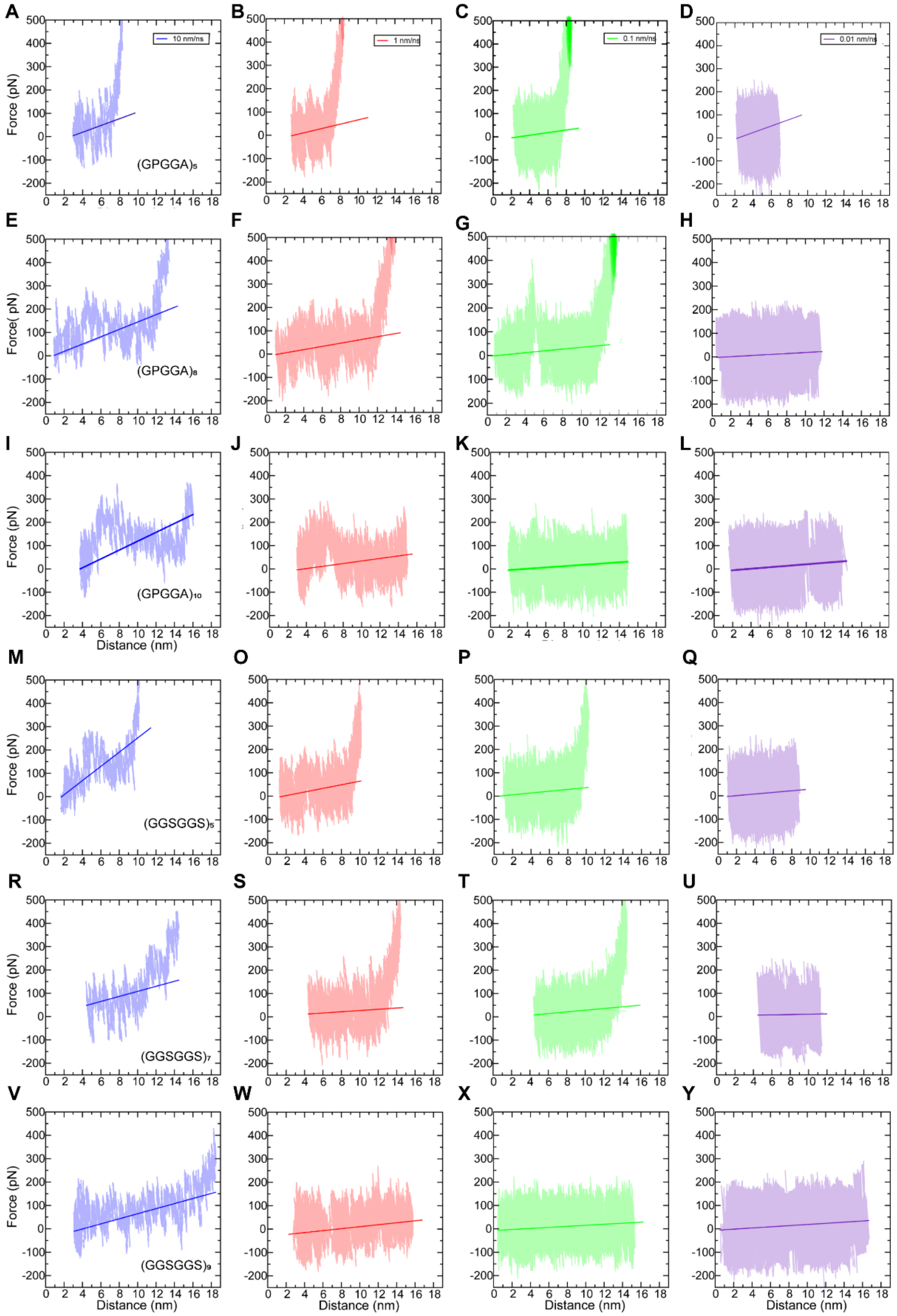


**Figure S4. Linear regressions of constant-velocity SMD data from 10 ns starting conformation.** (*A–L*) Linear regressions were used to obtain slope spring constants at 80% percent extension for $(GPGGA)_n$ linkers at different stretching velocities (0.01 nm/ns, 0.1 nm/ns, 1 nm/ns, and 10 nm/ns). (*M–Y*) Similar linear regressions for $(GGSGGS)_n$ systems.

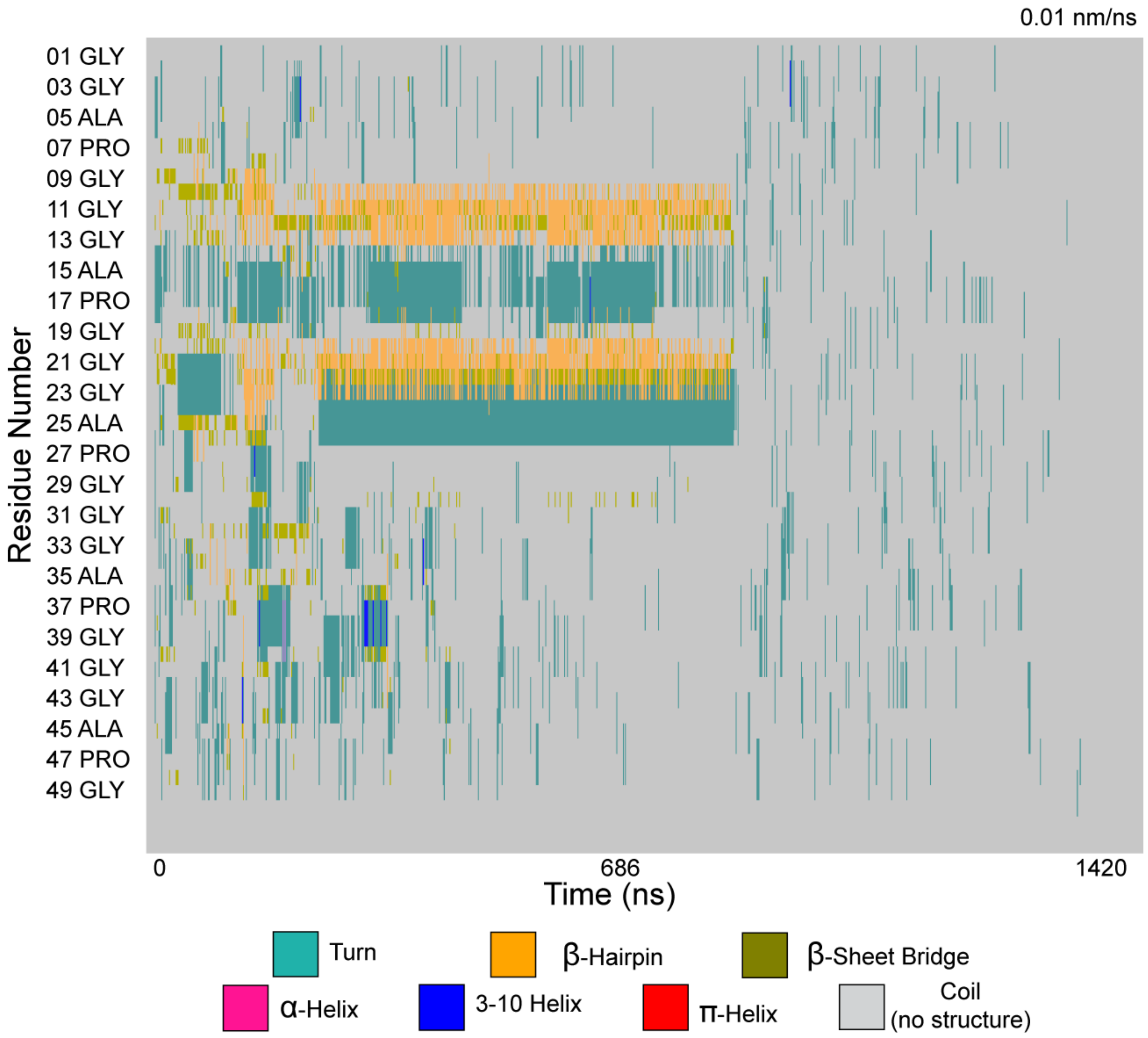


**Figure S5. Secondary structure analysis.** Secondary structure of spider silk linker $(GPGGA)_{10}$ stretched at 0.01 nm/ns shown per residue as a function of time.

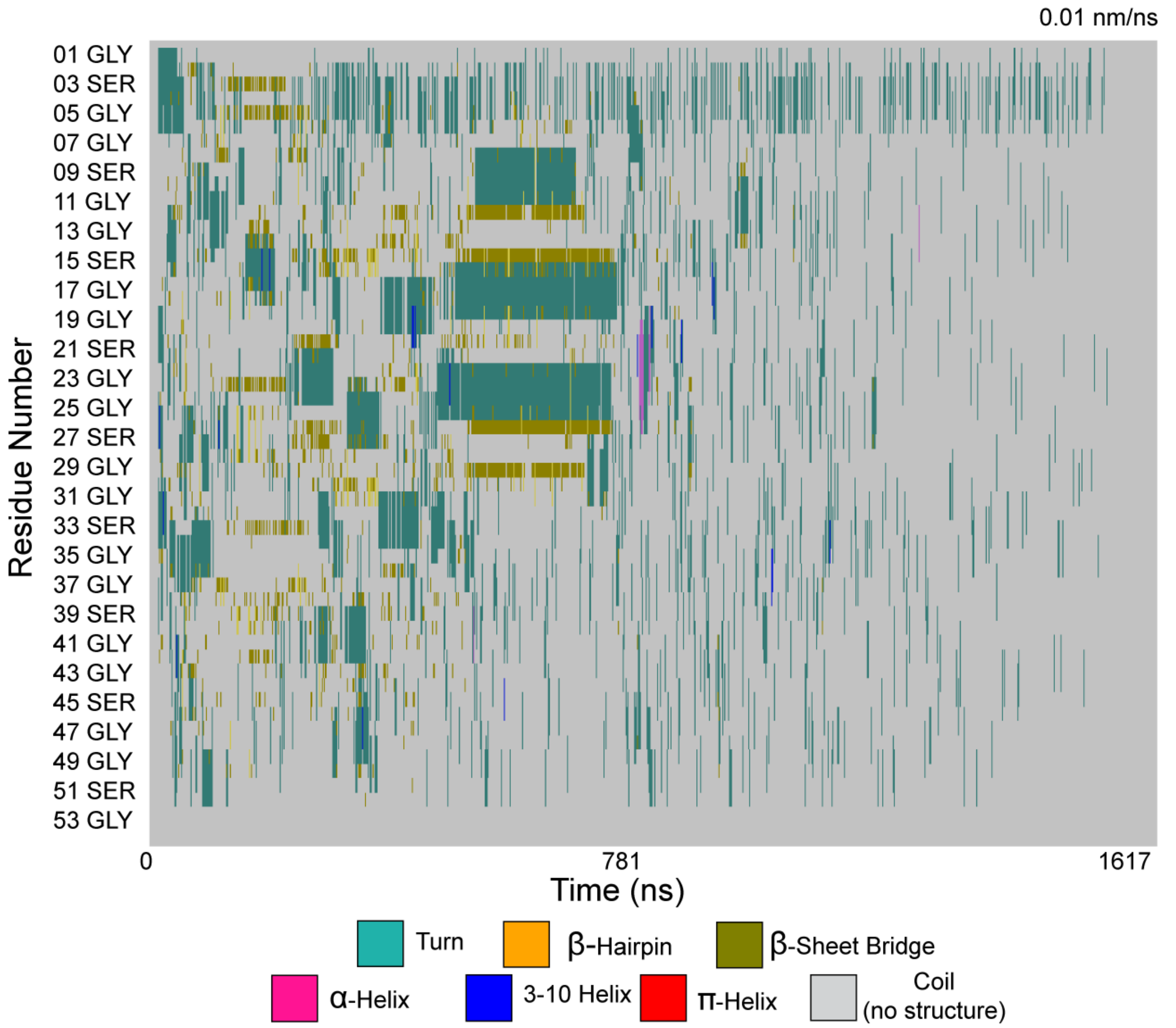


**Figure S6. Secondary structure analysis.** Secondary structure of synthetic linker $(GGSGGS)_9$ stretched at 0.01 nm/ns shown per residue as a function of time.

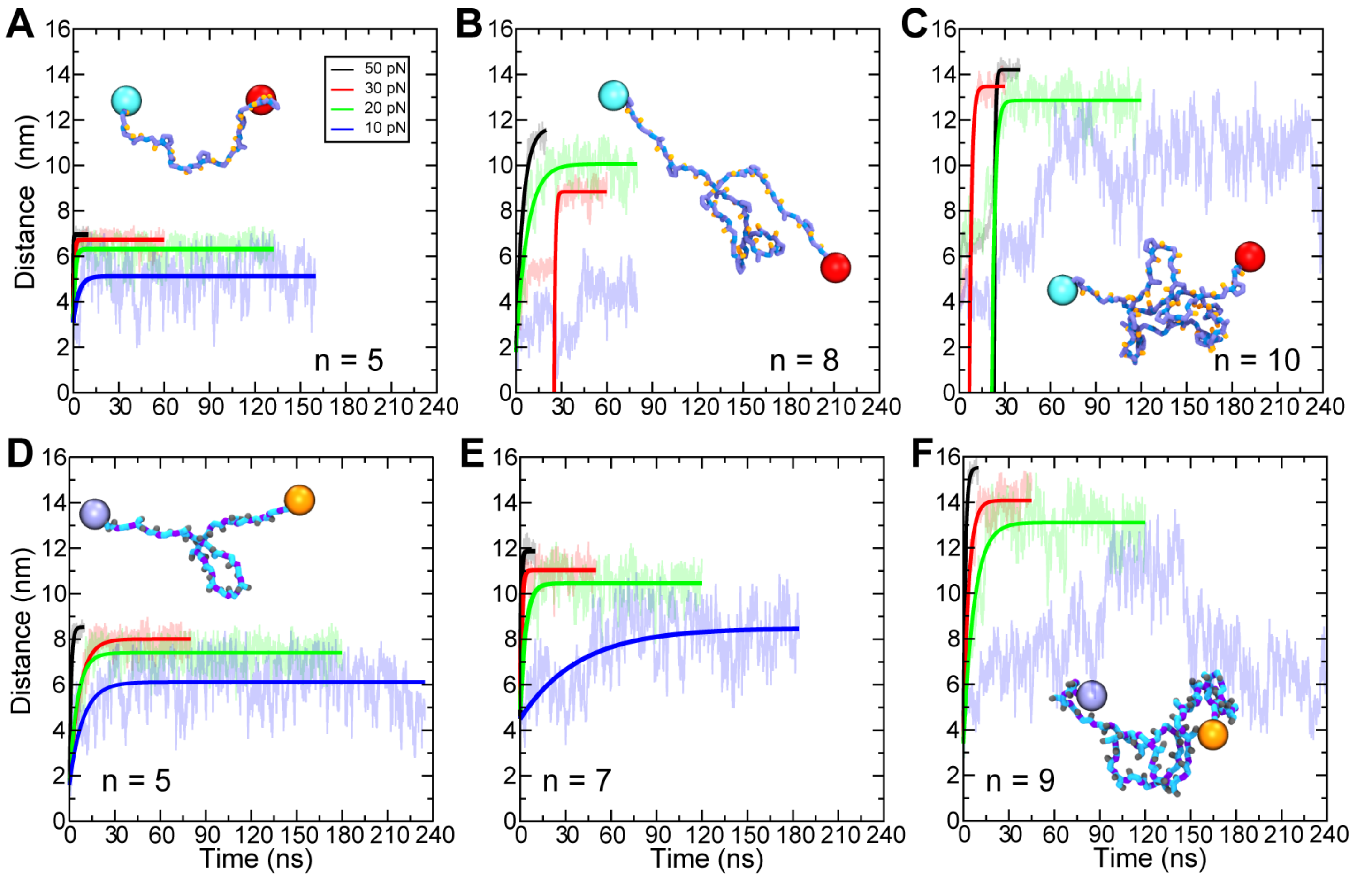


**Figure S7. Elasticity of spider silk $(GPGGA)_n$ and synthetic $(GGSGGS)_n$ sensors from constant-force stretching.** (*A–F*) Distance-time curves of SMD simulations performed at 10 pN (blue), 20 pN (green), 30 pN (red), and 50 pN (black) for $(GPGGA)_5$ (*A*), $(GPGGA)_8$ (*B*), $(GPGGA)_{10}$ (*C*), $(GGSGGS)_5$ (*D*), $(GGSGGS)_7$ (*E*), and $(GGSGGS)_9$ (*F*). Solid lines are fits used to obtain predicted spring constants. Insets show representative trajectory snapshots of each linker stretched at 20 pN as in Fig. 1 *B*.

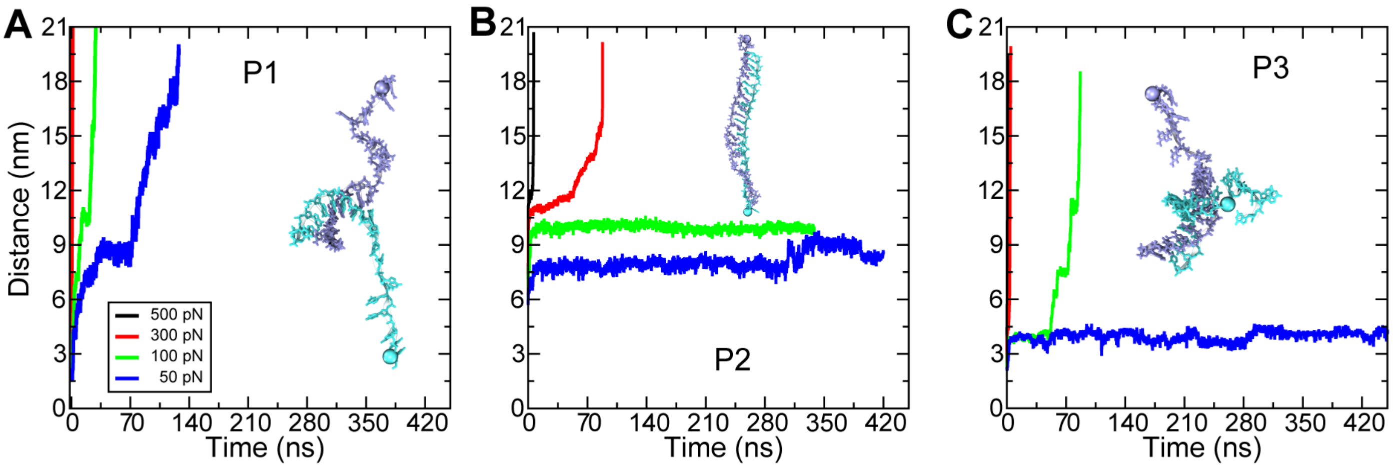


**Figure S8. Intermediate states in constant-force stretching of DNA-based sensors.** (*A–C*) Extension-time curves of DNA-based sensors stretched from different locations (P1, P2, and P3) at constant force (500 pN black; 300 pN red; 100 pN green; and 50 pN blue).

A

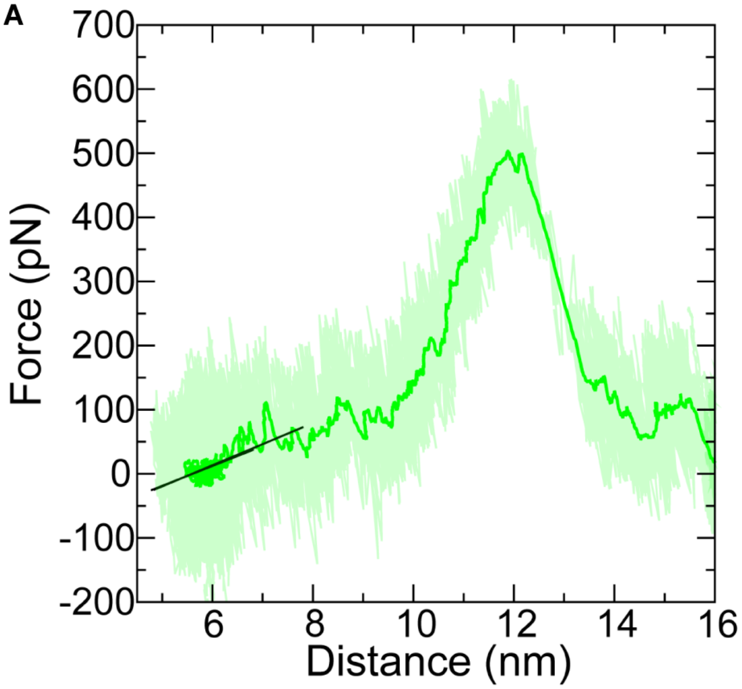


B

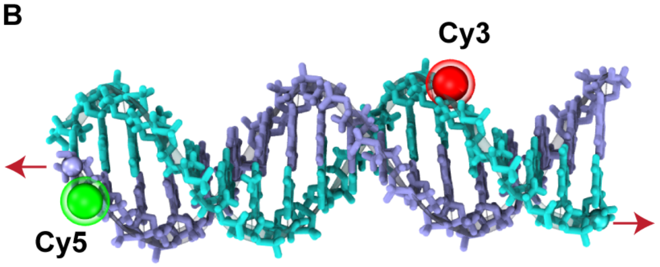


C

| Cy5-Cy3<br>$R_0$ 5.4 nm | | |
|---|---|---|
| FRET efficiency | 20 % - 80 % | 10 % - 90 % |
| Distance (nm) | 4.3 - 7 | 3.7 - 7.8 |
| Spring *k* (pN/nm) | 31.0 | 33.1 |

**Figure S9. Stretching DNA-based sensor.** (*A*) Force-extension curve of DNA-based sensor stretched from P3 at 0.1 nm/ns and a linear fit performed in the area of linear stretching. (*B*) Molecular model of a 18 dsDNA sensor with potential cy5-cy3 locations that could be used to detect linear stretching. (*C*) Summary table of FRET efficiency, stiffness and distance.

**Table S1.** Summary of SMD simulations of spider silk linker systems $(GPGGA)_n$.

| ***Label*** | **System** | $t_{sim}$ **(ns)** | **Type** | **Start** | **Velocity (nm/ns)** | **Force (pN)** | **Spring Constant ($k$)$^+$ (pN/nm)** | **Size (#atoms)** |
|---|---|---|---|---|---|---|---|---|
| *Sim1a* | | 15 | EQ | – | – | – | – | |
| *Sim1b* | | 2 | SMD | *Sim1a* | 10 | – | 14.9 | |
| *Sim1c* | | 10 | SMD | *Sim1a* | 1 | – | 5.2 | |
| *Sim1d* | | 155.6 | SMD | *Sim1a* | 0.1 | – | 3.7 | |
| *Sim1f* | | 10 | SMD | *Sim1a* | – | 50 | 10.5 | |
| *Sim1g* | | 30 | SMD | *Sim1a* | – | 30 | 6.3 | |
| *Sim1h* | | 58.3 | SMD | *Sim1a* | – | 20 | 4.2 | |
| *Sim1i* | **n = 5** | 160 | SMD | *Sim1a* | – | 10 | 2.5 | 71,446 |
| *Sim1j* | | 2 | SMD | *Sim1a** | 10 | – | 15.1 | |
| *Sim1k* | | 40 | SMD | *Sim1a** | 1 | – | 9.4 | |
| *Sim1l* | | 155.5 | SMD | *Sim1a** | 0.1 | – | 5.7 | |
| *Sim1m* | | 423 | SMD | *Sim1a** | 0.01 | – | 4.1 | |
| *Sim1n* | | 10 | SMD | *Sim1a** | – | 50 | 13.0 | |
| *Sim1o* | | 60 | SMD | *Sim1a** | – | 30 | 8.3 | |
| *Sim1p* | | 132.4 | SMD | *Sim1a** | – | 20 | 6.2 | |
| *Sim1q* | | 100 | SMD | *Sim1a** | – | 10 | 4.9 | |
| *Sim1a* | | 15 | EQ | – | – | – | – | |
| *Sim2b* | | 4 | SMD | *Sim2a* | 10 | – | 47.3 | |
| *Sim2c* | | 10 | SMD | *Sim2a* | 1 | – | 10.6 | |
| *Sim2d* | | 80 | SMD | *Sim2a* | 0.1 | – | 19.9 | |
| *Sim2f* | | 30 | SMD | *Sim2a* | – | 50 | 5.1 | |
| *Sim2g* | | 80 | SMD | *Sim2a* | – | 30 | 4.5 | |
| *Sim2h* | | 80 | SMD | *Sim2a* | – | 20 | N/A | |
| *Sim2i* | **n = 8** | 80 | SMD | *S1a* | – | 10 | N/A | 71,347 |
| *Sim2j* | | 1.7 | SMD | *Sim2a** | 10 | – | 16.3 | |
| *Sim2k* | | 17.1 | SMD | *Sim2a** | 1 | – | 6.9 | |
| *Sim2l* | | 165.7 | SMD | *Sim2a** | 0.1 | – | 4.3 | |
| *Sim2m* | | 1062.1 | SMD | *Sim2a** | 0.01 | – | 2.1 | |
| *Sim2n* | | 20 | SMD | *Sim2a** | – | 50 | 5.1 | |
| *Sim2o* | | 60 | SMD | *Sim2a** | – | 30 | 8.5 | |
| *Sim2p* | | 80 | SMD | *Sim2a** | – | 20 | 2.4 | |
| *Sim2q* | | 80 | SMD | *Sim2a** | – | 10 | N/A | |
| *Sim3a* | | 15 | EQ | – | – | – | – | |
| *Sim3b* | | 2 | SMD | *Sim3a* | 10 | – | 17.5 | |
| *Sim3c* | | 43.7 | SMD | *Sim3a* | 1 | – | 8.4 | |
| *Sim3d* | | 155.5 | SMD | *Sim3a* | 0.1 | – | 3.5 | |
| *Sim3f* | | 40 | SMD | *Sim3a* | – | 50 | 11.6 | |
| *Sim3g* | | 45 | SMD | *Sim3a* | – | 30 | 6.0 | 71,254 |
| *Sim3h* | | 90 | SMD | *Sim3a* | – | 20 | 3.7 | |
| *Sim3i* | **n = 10** | 240 | SMD | *Sim3a* | – | 10 | N/A | |
| *Sim3j* | | 1.5 | SMD | *Sim3a** | 10 | – | 19 | |
| *Sim3k* | | 13.3 | SMD | *Sim3a** | 1 | – | 5.2 | |
| *Sim3l* | | 132.5 | SMD | *Sim3a** | 0.1 | – | 2.8 | |
| *Sim3m* | | 1182.2 | SMD | *Sim3a** | 0.01 | – | 3.2 | |

| | | | | | | |
|---|---|---|---|---|---|---|
| *Sim3n* | 40 | SMD | *Sim3a** | – | 50 | 9.4 |
| *Sim3o* | 30 | SMD | *Sim3a** | – | 30 | 4.9 |
| *Sim3p* | 120 | SMD | *Sim3a** | – | 20 | 3.7 |
| *Sim3q* | 320 | SMD | S3a* | – | 10 | N/A |

[a]EQ indicates simulations that consisted of 1,000 steps of minimization, 1 ns of free dynamics in the *NpT* ensemble ($\gamma$ = 1 $ps^{-1}$), and 15 ns of free dynamics in the *NpT* ensemble ($\gamma$ = 0.1 $ps^{-1}$).

[*]Starting point after 10 ns of free dynamics in the *NpT* ensemble ($\gamma$ = 0.1 $ps^{-1}$).

[+]Spring constant calculated at 80% extension. Total simulation time = 5,700.1 ns

**Table S2.** Summary of SMD simulations for synthetic linker systems $(GGSGGS)_n$.

| ***Label*** | **System** | $t_{sim}$ **(ns)** | **Type** | **Start** | **Velocity (nm/ns)** | **Force (pN)** | **Spring Constant ($k$)$^+$** | **Size (#atoms)** |
|---|---|---|---|---|---|---|---|---|
| *Sim1a* | | 15 | EQ | – | – | – | – | |
| *Sim1b* | | 2 | SMD | *Sim1a* | 10 | – | 16.5 | |
| *Sim1c* | | 10 | SMD | *Sim1a* | 1 | – | 9.9 | |
| *Sim1d* | | 137 | SMD | *Sim1a* | 0.1 | – | 2.2 | |
| *Sim1f* | | 10 | SMD | *Sim1a* | – | 50 | 7.8 | |
| *Sim1g* | | 80 | SMD | *Sim1a* | – | 30 | 4.9 | |
| *Sim1h* | | 150 | SMD | *Sim1a* | – | 20 | 3.1 | 71,513 |
| *Sim1i* | **n = 5** | 235 | SMD | *Sim1a* | – | 10 | 2.2 | |
| *Sim1j* | | 2 | SMD | *Sim1a** | 10 | – | 30.5 | |
| *Sim1k* | | 10 | SMD | *Sim1a** | 1 | – | 7.7 | |
| *Sim1l* | | 100 | SMD | *Sim1a** | 0.1 | – | 4.4 | |
| *Sim1m* | | 750 | SMD | *Sim1a** | 0.01 | – | 3.7 | |
| *Sim1n* | | 10 | SMD | *Sim1a** | – | 50 | 7.2 | |
| *Sim1o* | | 80 | SMD | *Sim1a** | – | 30 | 4.6 | |
| *Sim1p* | | 180 | SMD | *Sim1a** | – | 20 | 3.4 | |
| *Sim1q* | | 234 | SMD | *Sim1a** | – | 10 | 2.2 | |
| *Sim1a* | | 15 | EQ | – | – | – | – | |
| *Sim2b* | | 4 | SMD | *Sim2a* | 10 | – | 15.2 | |
| *Sim2c* | | 30 | SMD | *Sim2a* | 1 | – | 6.7 | |
| *Sim2d* | | 164 | SMD | *Sim2a* | 0.1 | – | 4.3 | |
| *Sim2f* | | 15 | SMD | *Sim2a* | – | 50 | 5.5 | |
| *Sim2g* | | 50 | SMD | *Sim2a* | – | 30 | 3.4 | |
| *Sim2h* | **n = 7** | 200 | SMD | *Sim2a* | – | 20 | 2.6 | |
| *Sim2i* | | 284 | SMD | *Sim2a* | – | 10 | 2.2 | 71,244 |
| *Sim2j* | | 1.1 | SMD | *Sim2a** | 10 | – | 15.4 | |
| *Sim2k* | | 11 | SMD | *Sim2a** | 1 | – | 7.3 | |
| *Sim2l* | | 109 | SMD | *Sim2a** | 0.1 | – | 6.4 | |
| *Sim2m* | | 640 | SMD | *Sim2a** | 0.01 | – | 6 | |
| *Sim2n* | | 10 | SMD | *Sim2a** | – | 50 | 6.8 | |
| *Sim2o* | | 50 | SMD | *Sim2a** | – | 30 | 4.9 | |
| *Sim2p* | | 120 | SMD | *Sim2a** | – | 20 | 3.6 | |
| *Sim2q* | | 184 | SMD | *Sim2a** | – | 10 | 2.5 | |
| *Sim3a* | | 15 | EQ | – | – | – | – | |
| *Sim3b* | | 20.4 | SMD | *Sim3a* | 10 | – | 18.7 | |
| *Sim3c* | | 40 | SMD | *Sim3a* | 1 | – | 3.1 | |
| *Sim3d* | | 164.3 | SMD | *Sim3a* | 0.1 | – | 1.8 | |
| *Sim3f* | | 6.2 | SMD | *Sim3a* | – | 50 | 2.5 | |
| *Sim3g* | | 33 | SMD | *Sim3a* | – | 30 | 2.0 | 71,206 |
| *Sim3h* | | 200 | SMD | *Sim3a* | – | 20 | 1.9 | |
| *Sim3i* | **n = 9** | 384 | SMD | *Sim3a* | – | 10 | N/A | |
| *Sim3j* | | 1.9 | SMD | *Sim3a** | 10 | – | 10.2 | |
| *Sim3k* | | 15 | SMD | *Sim3a** | 1 | – | 4.2 | |
| *Sim3l* | | 147.4 | SMD | *Sim3a** | 0.1 | – | 2.4 | |
| *Sim3m* | | 1617.7 | SMD | *Sim3a** | 0.01 | – | 1.9 | |
| *Sim3n* | | 10 | SMD | *Sim3a** | – | 50 | 4.1 | |
| *Sim3o* | | 45 | SMD | *Sim3a** | – | 30 | 2.8 | |

| | | | | | | |
|---|---|---|---|---|---|---|
| *Sim3p* | 120 | SMD | *Sim3a** | – | 20 | 2.1 |
| *Sim3q* | 284 | SMD | *Sim3a** | – | 10 | N/A |

[a]EQ indicates simulations that consisted of 1,000 steps of minimization, 1 ns of free dynamics in the *NpT* ensemble ($\gamma = 1$ $ps^{-1}$), and 15 ns of free dynamics in the *NpT* ensemble ($\gamma = 0.1$ $ps^{-1}$).

[*]Starting point after 10 ns of free dynamics in the *NpT* ensemble ($\gamma = 0.1$ $ps^{-1}$).

[+]Spring constant calculated at 80% extension. Total simulation time = 6,996 ns

**Table S3.** Summary of percentage of secondary structure motifs during 0.01 nm/ns stretching simulations.

| System | Motif | | | | | | |
|---|---|---|---|---|---|---|---|
| | T (%) | E (%) | B (%) | H (%) | G (%) | I (%) | C (%) |
| **(GPGGA)$_5$** | 10.68 | 0.15 | 0.32 | 0.00 | 0.12 | 0.00 | 88.73 |
| **(GPGGA)$_8$** | 13.60 | 0.64 | 2.00 | 0.02 | 0.92 | 0.00 | 82.82 |
| **(GPGGA)$_{10}$** | 12.06 | 4.58 | 2.05 | 0.01 | 0.10 | 0.00 | 81.19 |
| **(GGSGGS)$_5$** | 14.78 | 0.56 | 1.66 | 0.00 | 0.32 | 0.01 | 82.67 |
| **(GGSGGS)$_7$** | 12.67 | 0.01 | 0.29 | 0.04 | 0.22 | 0.01 | 86.75 |
| **(GGSGGS)$_9$** | 14.70 | 0.46 | 3.13 | 0.11 | 0.09 | 0.00 | 81.50 |

**T** = Turn **E** = β Hairpin **B** = β sheet bridge **H** = α helix **G** = 3-10 helix **I** = π-helix **C** = Coil (no structure)

**Table S4.** Spring constants at different stretching forces.

| *System* | Simulation Pulling Force | Spring Constant *k* (pN/nm)* | Friction coefficient $\gamma_f$ (pN·ns/nm) | Spring Constant *k* (pN/nm)** | Friction coefficient $\gamma_f$ (pN·ns/nm) |
|---|---|---|---|---|---|
| | 50 | 13.0 | 2.3 | 10.5 | 5.7 |
| | 30 | 8.3 | 5.8 | 6.3 | 6.5 |
| **$(GPGGA)_5$** | 20 | 6.2 | 7.6 | 4.2 | 7.1 |
| | 10 | 4.9 | 17.6 | 2.5 | 95.3 |
| | 50 | 5.1 | 23.4 | 5.1 | 9.3 |
| | 30 | 8.5 | 6.7 | 4.5 | 10.3 |
| **$(GPGGA)_8$** | 20 | 2.4 | 17.8 | N/A | N/A |
| | 10 | N/A | N/A | N/A | N/A |
| | 50 | 9.4 | 6.1 | 11.6 | 6.8 |
| | 30 | 4.9 | 6.7 | 6.0 | 8.6 |
| **$(GPGGA)_{10}$** | 20 | 3.7 | 7.1 | 3.7 | 10.0 |
| | 10 | N/A | N/A | N/A | N/A |
| | 50 | 7.2 | 7.9 | 7.8 | 1.75 |
| | 30 | 4.6 | 30.1 | 4.9 | 2.7 |
| **$(GGSGGS)_5$** | 20 | 3.4 | 18.1 | 3.1 | 74.9 |
| | 10 | 2.2 | 19.4 | 2.2 | 1.6 |
| | 50 | 6.8 | 4.2 | 5.5 | 7.8 |
| | 30 | 4.9 | 5.2 | 3.4 | 6.4 |
| **$(GGSGGS)_7$** | 20 | 3.6 | 13.2 | 2.6 | 13.7 |
| | 10 | 2.5 | 95.8 | 2.2 | 22.2 |
| | 50 | 4.1 | 5.3 | 2.5 | 18.0 |
| | 30 | 2.8 | 9.4 | 2.0 | 30.0 |
| **$(GGSGGS)_9$** | 20 | 2.1 | 14.5 | 1.9 | 21.0 |
| | 10 | N/A | N/A | N/A | N/A |

*Starting point after 10 ns of free dynamics *NpT* ensemble ($\gamma = 0.1\ ps^{-1}$).

**Starting point after 15 ns of free dynamics *NpT* ensemble ($\gamma = 0.1\ ps^{-1}$).

**Table S5.** Summary of SMD simulations for DNA systems.

| ***Label*** | **Stretching Configuration** | **$t_{sim}$ (ns)** | **Type** | **Start** | **Velocity (nm/ns)** | **Stretching Force (pN)** | **Force Peak (pN)** | **Size (#atoms)** |
|---|---|---|---|---|---|---|---|---|
| *Sim1a* | | 10 | EQ | – | – | – | – | |
| *Sim1b* | | 2.3 | SMD | *Sim1a* | 10 | – | 468 | |
| *Sim1c* | | 20.2 | SMD | *Sim1a* | 1 | – | 191 | |
| *Sim1d* | P1 | 166 | SMD | *Sim1a* | 0.1 | – | 163 | 214,550 |
| *Sim1e* | | 0.8 | SMD | *Sim1a* | – | 500 | – | |
| *Sim1f* | | 1.6 | SMD | *Sim1a* | – | 300 | – | |
| *Sim1g* | | 29 | SMD | *Sim1a* | – | 100 | – | |
| *Sim1h* | | 127.7 | SMD | *Sim1a* | – | 50 | – | |
| *Sim2b* | | 2.1 | SMD | *Sim1a* | 10 | – | 652 | |
| *Sim2c* | | 27.8 | SMD | *Sim1a* | 1 | – | 274 | |
| *Sim2d* | | 152.1 | SMD | *Sim1a* | 0.1 | – | 217 | |
| *Sim2e* | P3 | 0.9 | SMD | *Sim1a* | – | 500 | – | 214,550 |
| *Sim2f* | | 4.5 | SMD | *Sim1a* | – | 300 | – | |
| *Sim2g* | | 87 | SMD | *Sim1a* | – | 100 | – | |
| *Sim2h* | | 509.6 | SMD | *Sim1a* | – | 50 | – | |
| *Sim3b* | | 1.9 | SMD | *Sim1a* | 10 | – | 1,242 | |
| *Sim3c* | | 15.5 | SMD | *Sim1a* | 1 | – | 752 | |
| *Sim3d* | | 139.2 | SMD | *Sim1a* | 0.1 | – | 490 | |
| *Sim3e* | P2 | 6.6 | SMD | *Sim1a* | – | 500 | – | 214,550 |
| *Sim3f* | | 87.9 | SMD | *Sim1a* | – | 300 | – | |
| *Sim3g* | | 338.4 | SMD | *Sim1a* | – | 100 | – | |
| *Sim3h* | | 420.2 | SMD | *Sim1a* | – | 50 | – | |

[a]EQ indicates simulations that consisted of 1,000 steps of minimization, 1 ns of free dynamics in the *NpT* ensemble ($\gamma = 1$ ps$^{-1}$), and 15 ns of free dynamics in the *NpT* ensemble ($\gamma = 0.1$ ps$^{-1}$).

Total simulation time = 2,115 ns.

**Table S6.** Spring constants at different stretching velocities using 80% extension.

| System | Velocity | Spring Constant *k* (pN/nm)* | Spring Constant *k* (pN/nm)** |
|---|---|---|---|
| **(GPGGA)$_5$** | 10 | 15.1 | 14.9 |
| | 1 | 9.4 | 5.2 |
| | 0.1 | 5.7 | 3.7 |
| | 0.01 | 4.1 | - |
| **(GPGGA)$_8$** | 10 | 16.3 | 47.3 |
| | 1 | 6.9 | 10.6 |
| | 0.1 | 4.3 | 19.9 |
| | 0.01 | 2.1 | - |
| **(GPGGA)$_{10}$** | 10 | 19 | 17.5 |
| | 1 | 5.2 | 8.4 |
| | 0.1 | 2.8 | 3.5 |
| | 0.01 | 3.2 | - |
| **(GGSGGS)$_5$** | 10 | 30.5 | 16.5 |
| | 1 | 7.7 | 9.9 |
| | 0.1 | 4.4 | 2.2 |
| | 0.01 | 3.7 | - |
| **(GGSGGS)$_7$** | 10 | 15.4 | 15.2 |
| | 1 | 7.3 | 6.7 |
| | 0.1 | 6.4 | 4.3 |
| | 0.01 | 6 | - |
| **(GGSGGS)$_9$** | 10 | 10.2 | 18.7 |
| | 1 | 4.2 | 3.1 |
| | 0.1 | 2.4 | 1.8 |
| | 0.01 | 1.9 | - |

*Starting point after 10 ns of free dynamics in the *NpT* ensemble ($\gamma = 0.1$ ps$^{-1}$).
**Starting point after 15 ns of free dynamics in the *NpT* ensemble ($\gamma = 0.1$ ps$^{-1}$).

**Table S7.** Average spring constants across all constant-velocity simulations of $(GPGGA)_n$ and $(GGSGGS)_n$ calculated for extensions of selected FRET pairs after 10 ns equilibration.

| System | Velocity (nm/ns) | Clover-mRuby2 *k* (pN/nm) | mTurquoise2-sEYFP *k* (pN/nm) | LSSmOrange-mKate2 *k* (pN/nm) | ECFP-EYFP *k* (pN/nm) |
|---|---|---|---|---|---|
| **$(GPGGA)_5$** | 10 | 12.8 | 21.1 | 14.9 | 28.1 |
| | 1 | 7.4 | 7.3 | 7.8 | -1.7 |
| | 0.1 | 3.7 | 3.3 | 4.2 | 3.7 |
| | 0.01 | 4.5 | 4.4 | 4.5 | 5.3 |
| **$(GPGGA)_8$** | 10 | 40.9 | 44.1 | 36.2 | 46 |
| | 1 | 15.9 | 19.1 | 10.6 | 18 |
| | 0.1 | 23.4 | 26.2 | 20.5 | 13.6 |
| | 0.01 | 1.9 | 2.3 | 1.8 | 2.9 |
| **$(GPGGA)_{10}$** | 10 | 79.2 | 62.7 | 81.9 | 44.9 |
| | 1 | 62.6 | 75.6 | 62.7 | 71.2 |
| | 0.1 | 25.9 | 29.7 | 16.3 | -2.06 |
| | 0.01 | 4.1 | 3.0 | 4.1 | -5.0 |
| **$(GGSGGS)_5$** | 10 | 49.3 | 52.5 | 47.3 | 48.3 |
| | 1 | 13.2 | 16.1 | 10.2 | 22.5 |
| | 0.1 | 5.7 | 6.7 | 4.9 | 11.4 |
| | 0.01 | 2.8 | 2.8 | 2.7 | 3.6 |
| **$(GGSGGS)_7$** | 10 | 14.7 | 14.1 | 15.1 | 28.7 |
| | 1 | 9.1 | 10.2 | 7.6 | 23.2 |
| | 0.1 | 6.8 | 7.5 | 6.4 | 17 |
| | 0.01 | 6.9 | 7.2 | 6.3 | 20.5 |
| **$(GGSGGS)_9$** | 10 | 32.9 | 35.3 | 25.1 | 56.9 |
| | 1 | 4.5 | 5.1 | 3.6 | 6.3 |
| | 0.1 | 7.2 | 3.9 | 6.2 | 3.8 |
| | 0.01 | 4.6 | 4.6 | 4.1 | -5.9 |

**Table S8.** Average spring constants across all constant-velocity simulations of $(GPGGA)_n$ and $(GGSGGS)_n$ calculated for extensions of selected FRET pairs after 15 ns equilibration.

| System | Velocity (nm/ns) | Clover-mRuby2 *k* (pN/nm) | mTurquoise2-sEYFP *k* (pN/nm) | LSSmOrange-mKate2 *k* (pN/nm) | ECFP-EYFP *k* (pN/nm) |
|---|---|---|---|---|---|
| **$(GPGGA)_5$** | 10 | 17.1 | 20.6 | 12 | 32.6 |
| | 1 | 4.5 | 6.6 | 2.8 | 9.7 |
| | 0.1 | 2.4 | 3 | 2.9 | 1.9 |
| **$(GPGGA)_8$** | 10 | 18.7 | 46.8 | 41.6 | 39.2 |
| | 1 | 4.1 | 11.2 | 10.6 | 15.8 |
| | 0.1 | 4.5 | 19.1 | 15.1 | 7.9 |
| **$(GPGGA)_{10}$** | 10 | 58.1 | 48.2 | 72.3 | 26.4 |
| | 1 | 43.1 | 38.1 | 58.2 | 30.1 |
| | 0.1 | 28.9 | 19.1 | 44.1 | 0.9 |
| **$(GGSGGS)_5$** | 10 | 14.1 | 11.8 | 16 | 12.5 |
| | 1 | 8.3 | 7.4 | 9.3 | 8.1 |
| | 0.1 | 2.8 | 3.9 | 2.5 | 6.1 |
| **$(GGSGGS)_7$** | 10 | 20.5 | 17.6 | 20.7 | 12 |
| | 1 | 8.4 | 8.1 | 7.8 | 6.9 |
| | 0.1 | 4.1 | 4.2 | 4.1 | 4 |
| **$(GGSGGS)_9$** | 10 | 32.7 | 33.5 | 34 | 41.8 |
| | 1 | 4.9 | 6.4 | 3.5 | 7 |
| | 0.1 | 4.2 | 4.2 | 3.4 | 5.7 |

**Table S9.** Range of extensions used for analysis of spring constants (39,63,64).

| System | 80% of extension (nm) | Clover-mRuby2 (nm) | mTurquoise2-sEYFP (nm) | LSSmOrange-mKate2 (nm) | ECFP-EYFP (nm) |
|---|---|---|---|---|---|
| **(GPGGA)$_5$** | 0 – 7.0 | | | | |
| **(GPGGA)$_8$** | 0 – 11.2 | | | | |
| **(GPGGA)$_{10}$** | 0 – 14.1 | 3.0 – 6.0 | 2.5 – 5.5 | 3.5 – 6.5 | 1.5 – 4.5 |
| **(GGSGGS)$_5$** | 0 – 8.4 | | | | |
| **(GGSGGS)$_7$** | 0 – 11.8 | | | | |
| **(GGSGGS)$_9$** | 0 – 15.1 | | | | |
| **Förster Radius** | | 6.3 | 5.9 | 7.0 | 4.9 |
